# The *Aquilegia* genome reveals a hybrid origin of core eudicots

**DOI:** 10.1101/407973

**Authors:** Gökçe Aköz, Magnus Nordborg

**Affiliations:** Gregor Mendel Institute, Austrian Academy of Sciences, Vienna Biocenter, Vienna, Austria; Vienna Graduate School of Population Genetics, Vienna, Austria

## Abstract

**Background:** Whole-genome duplications (WGD) have dominated the evolutionary history of plants. One consequence of WGD is a dramatic restructuring of the genome as it undergoes diploidization, a process under which deletions and rearrangements of various sizes scramble the genetic material, leading to a repacking of the genome and eventual return to diploidy. Here, we investigate the history of WGD in the columbine genus *Aquilegia*, a basal eudicot, and use it to illuminate the origins of the core eudicots.

**Results:** Within-genome synteny confirms that columbines are ancient tetraploids, and comparison with the grape genome reveals that this tetraploidy appears to be shared with the core eudicots. Thus, the ancient *gamma* hexaploidy found in all core eudicots must have involved a two-step process: first tetraploidy in the ancestry of all eudicots, then hexaploidy in the ancestry of core eudicots. Furthermore, the precise pattern of synteny sharing suggests that the latter involved allopolyploidization, and that core eudicots thus have a hybrid origin.

**Conclusions:** Novel analyses of synteny sharing together with the well-preserved structure of the columbine genome reveal that the *gamma* hexaploidy at the root of core eudicots is likely a result of hybridization between a tetraploid and a diploid species.

## Background

Whole-genome duplication (WGD) is common in the evolutionary history of plants [reviewed in 1,2]. All flowering plants are descended from a polyploid ancestor, which in turn shows evidence of an even older WGD shared by all seed plants [3]. These repeated cycles of polyploidy dramatically restructure plant genomes. Presumably driven by the “diploidization” process, whereby genomes are returned to an effectively diploid state, chromosomes are scrambled via fusions and fissions, lose both repetitive and genic sequences, or are lost entirely [4–11]. Intriguingly, gene loss after WGD is non-random: not only is there a bias against the retention of certain genes [12,13], but also against the retention of one of the WGD-derived paralog chromosomes [6,9,14–16].

We investigated the history of WGDs in the columbine genus *Aquilegia* for two reasons. The first is related to its phylogenetic position: columbines are “basal” eudicots, having diverged early from the remaining “core” eudicots [17,18]. This matters because our understanding of eudicot karyotype evolution is limited to the heavily sampled core eudicots. Using the recently published *Aquilegia coerulea* genome [19], we were able to address key questions about the history of polyploidization in all eudicots. Second, we traced the origins of the columbine chromosomes with a particular focus on the strange chromosome 4, which differs from the rest of the genome many ways. In particular, it harbors more genetic polymorphism and transposable elements, has lower gene density and reduced gene expression, and appears to migrate more, including between species. It also carries the rDNA clusters, and there is reason to believe that knowing the history of the chromosome could help explain its aberrant behavior [19].

## Results

### Within genome synteny confirms columbine paleotetraploidy

Ancient WGDs have been commonly inferred from the distribution of divergences between gene duplicates. The simultaneous generation of gene duplicates via WGD is expected to produce a peak in the age distribution relative to the background age distribution of single gene duplicates [20–22]. Such a spike of ancient gene birth was the first evidence of paleotetraploidy in columbines [23], and was later supported by gene count-based modelling [24].

Given an assembled genome, a more direct method to infer ancient polyploidy is to look for regions with conserved gene order [25,26]. Such conservation (a.k.a., synteny) decreases over time due to gene loss and rearrangements, but will still be detectable unless the extent of change is too extensive. We detected a total of 121 synteny block pairs harboring at least five paralogous gene pairs within the columbine genome. The distribution of these blocks across the seven columbine chromosomes indicates pairwise synteny between large genomic regions (Fig. 1). This 1:1 relationship suggests a single round of WGD in columbine, and is further supported by similar levels of divergence between synteny pairs (Figs. S1 and S2).

**Fig. 1:**
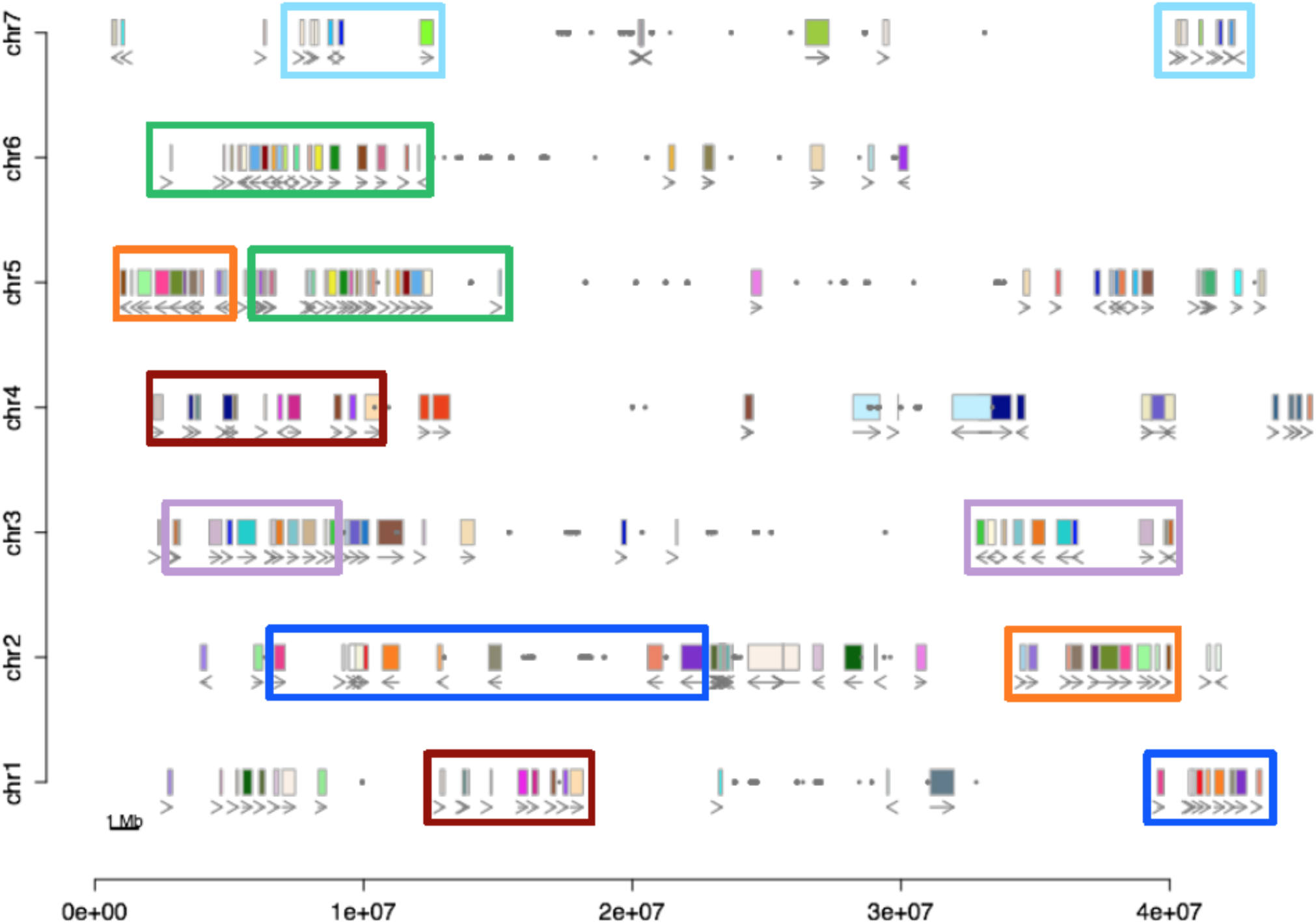
Intragenomic synteny blocks in the columbine genome. Pairs of synteny blocks are denoted as uniquely colored small rectangles. Larger rectangles of the same color outline large regions of synteny. Arrows under the synteny blocks show the orientation of the alignment between collinear genes. Grey dots highlight BLAST hits of a 329 bp centromeric repeat monomer [19,27].

### Columbines share ancient tetraploidy with core eudicots

All sequenced core eudicots appear to share a triplicate genome structure due to paleohexaploidy postdating the separation of monocots and eudicots [9,28–32, and Supplementary Note 5 in 33]. The tetraploidy in columbines, a basal eudicot, might be independent of this ancient “*gamma*” hexaploidy (Scenarios 1 and 2 in Fig. 2) or might be a remnant of a WGD at the base of all eudicots, which formed the first step of the *gamma* hexaploidy in core eudicots (Scenario 3 in Fig.2).

**Fig. 2:**
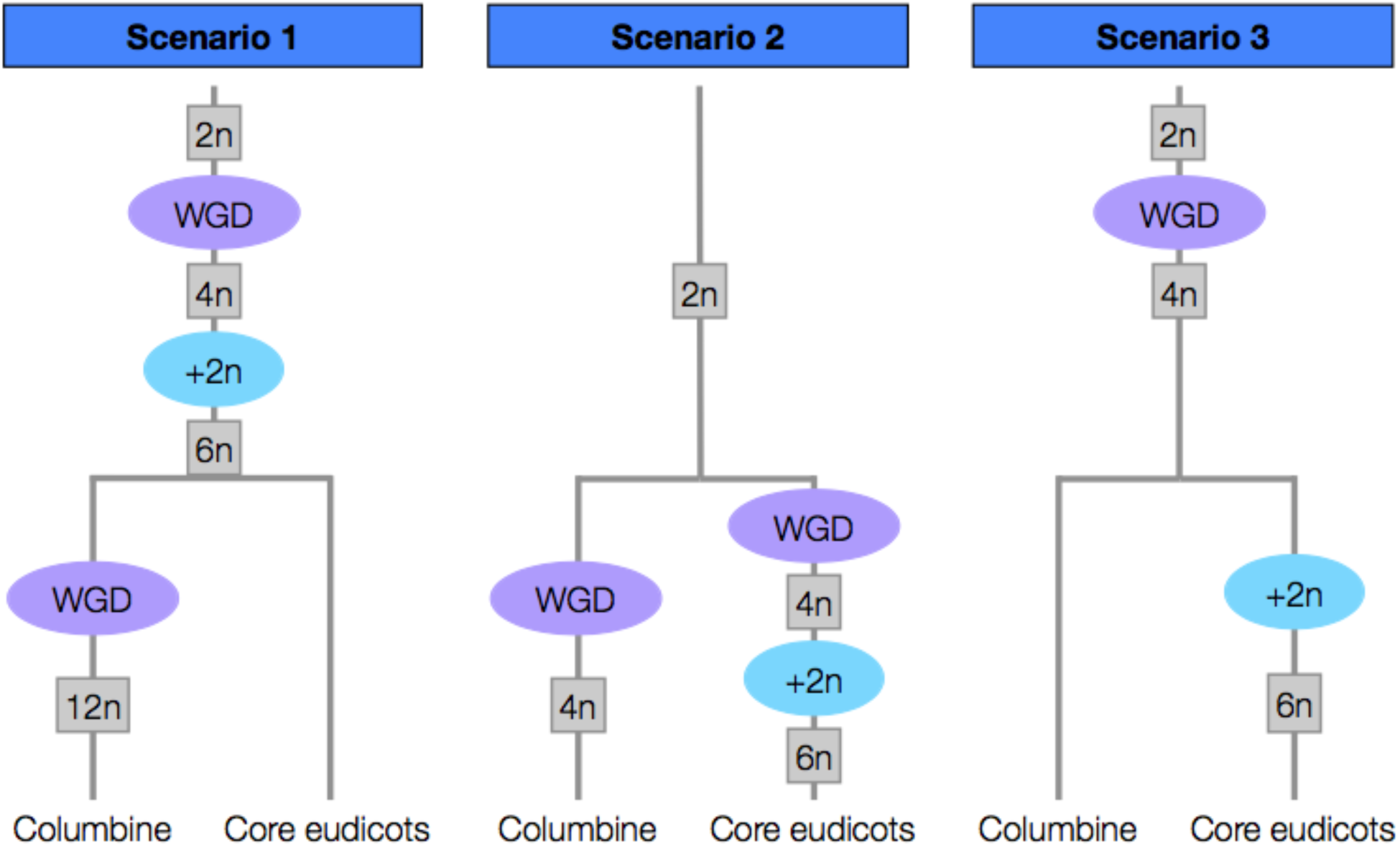
Three scenarios for the relationship between columbine tetraploidy and core eudicot “*gamma*” hexaploidy. The *gamma* hexaploidy is a two-step process: a single round of WGD creates tetraploids (4n) whose unreduced gametes then fuse with diploid gametes (+2n). **Scenario 1**: *gamma* hexaploidy precedes the split between columbine and core eudicots, with the former undergoing an additional tetraploidy. **Scenario 2**: Both *gamma* hexaploidy and columbine tetraploidy occur after the split between columbines and core eudicots. **Scenario 3**: Columbine tetraploidy is derived from the ancient tetraploidy that was the first step of the process leading to *gamma* hexaploidy.

We used the grape (*Vitis vinifera*) genome as a representative of the core eudicots to distinguish between the three scenarios in Fig. 2. Grape has experienced a relatively small number of chromosomal rearrangements post-*gamma* and thus strongly resembles the ancestral pre-hexaploid genome [34]. Given the ploidy level of columbine under each scenario, we can predict the ratio of haploid chromosome sets in grape to that in columbine. If tetraploidy in columbines is lineage-specific and superimposed on the *gamma* hexaploidy (Scenario 1), we would expect to find a 3:6 ratio of grape and columbine synteny blocks. Instead, we observe a 3:2 relationship (Figs. 3 and S3) as expected under Scenarios 2 or 3. A similar 3:2 pattern is found in comparisons between grape and sacred lotus [35]. This strongly suggests that basal eudicots do not share the triplicate genome structure of core eudicots, ruling out Scenario 1.

**Fig. 3:**
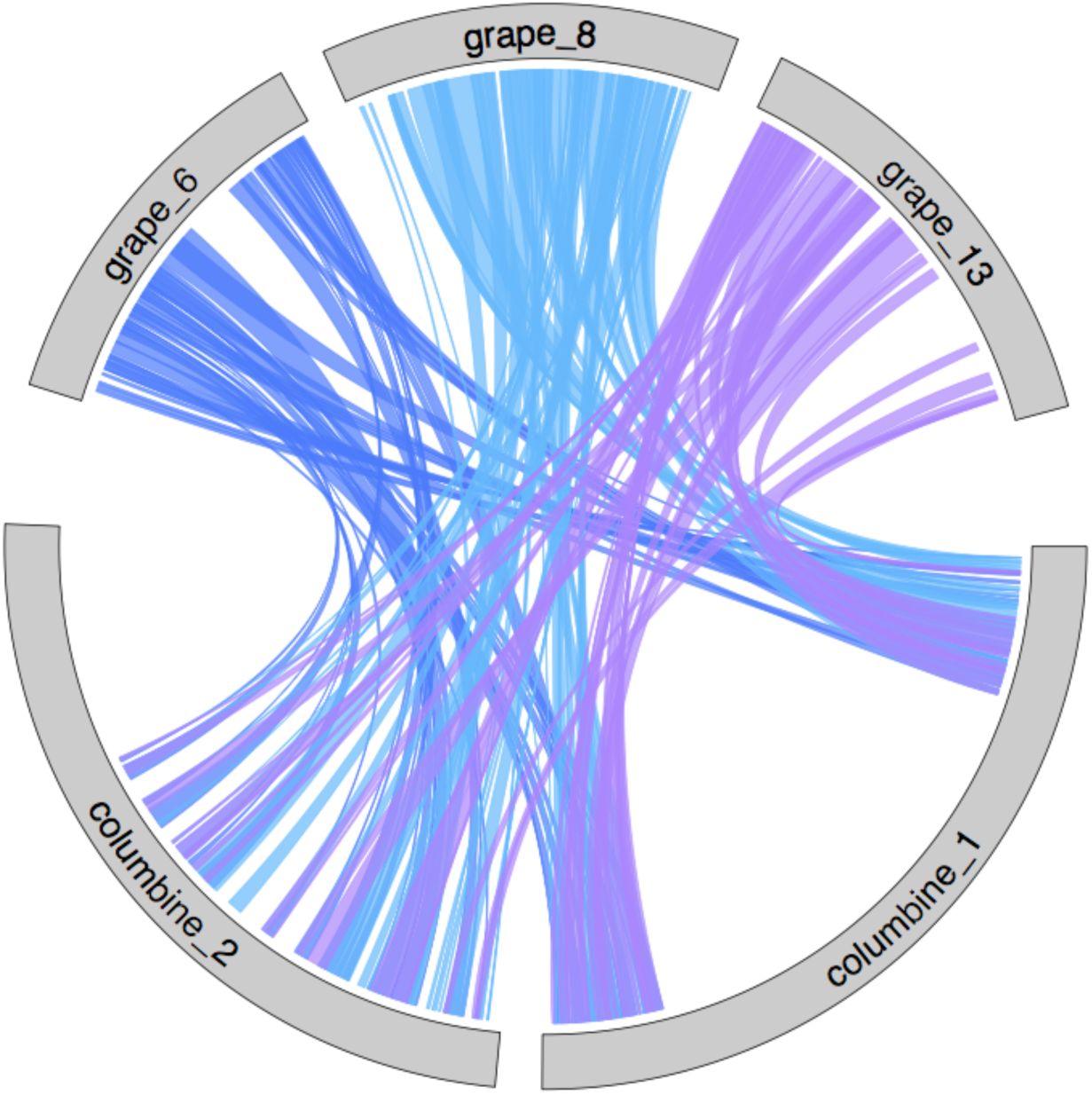
Synteny between the homologous regions of columbine and grape. The results for columbine chromosomes 1 and 2, and grape chromosomes 6, 8 and 13, shown here, reflect the overall synteny relationship of 3:2 between grape:columbine chromosomes (see Fig. S3 and Supplementary Data 2). This pattern argues against Scenario 1, but is consistent with either Scenario 2 or Scenario 3 in Fig. 2.

Distinguishing between the two remaining scenarios is more difficult. We began by comparing the divergence at synonymous sites (Ks) between columbine paralogs, grape paralogs and columbine-grape homologs. In agreement with the analysis of Jiao et al. [36], the Ks distribution for grape paralogs appears to two major peaks, as expected under the two-step model for *gamma* hexaploidy (Fig. 4). However, columbine paralogs and columbine-grape homologs each show a single peak of divergence — and the peaks overlap each other and the “older” divergence peak of grape paralogs. This suggests that columbine tetraploidy is derived from the tetraploidy that eventually led to *gamma* hexaploidy in core eudicots (Scenario 3).

**Fig. 4:**
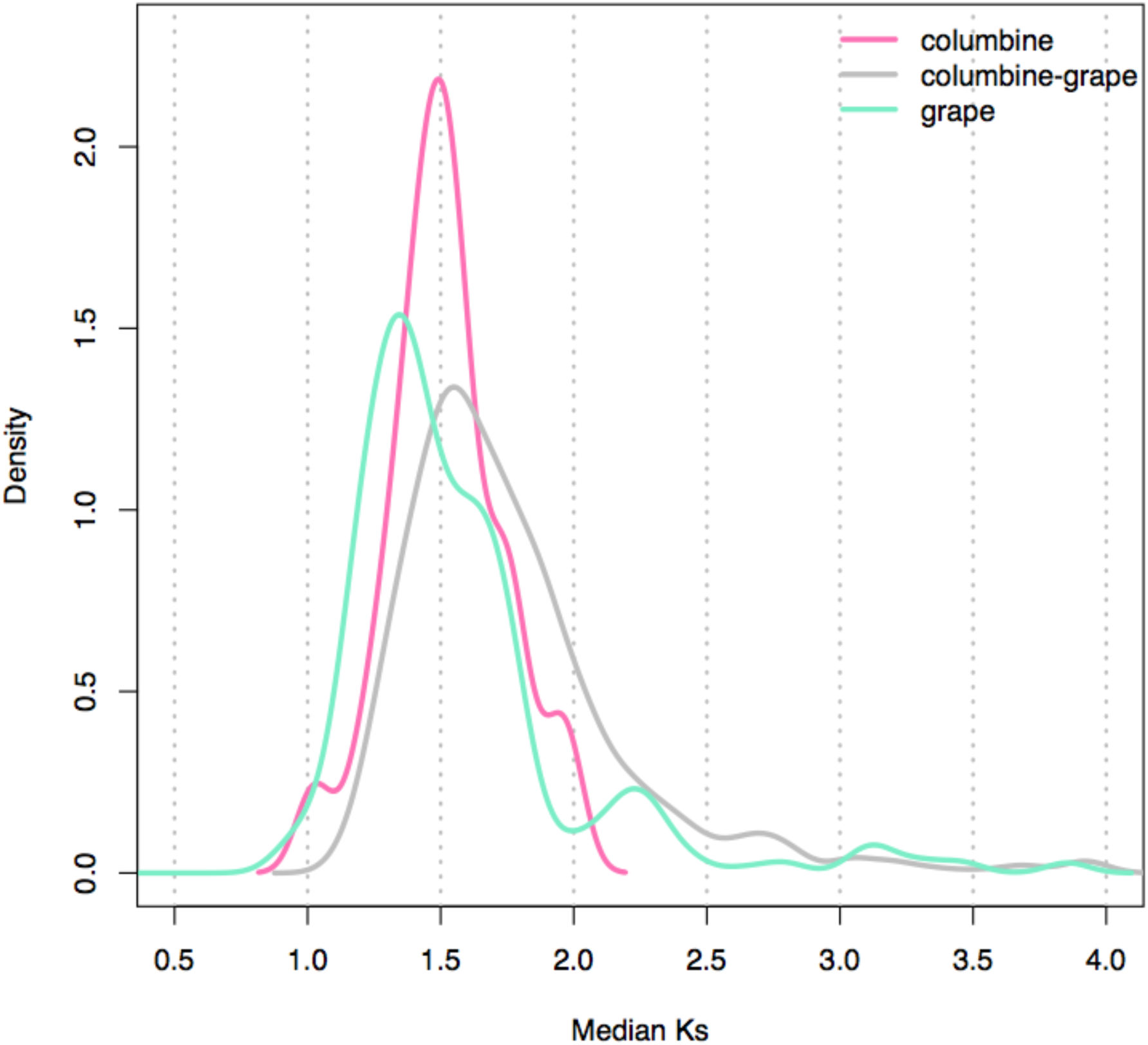
The distribution of the median Ks across syntenic regions. Synteny blocks are identified within columbine, between columbine and grape, and within grape. Note that only the putative WGD-derived blocks (median Ks=1-2) are kept in columbine (Fig. S2).

Next we considered the gene-order similarity between pairs of columbine and grape chromosomes. If columbine and grape have descended from a common tetraploid ancestor (Scenario 3), they should have inherited all diploidization-driven changes to gene order between the paralogous chromosomes of the tetraploid ancestor. As a result, we expect to see alternative paralogous gene orders to be uniquely shared between two pairs of columbine and grape chromosomes (Fig. 5) — whereas if tetraploidization occurred twice (Scenario 2), no such pattern is expected.

**Fig. 5:**
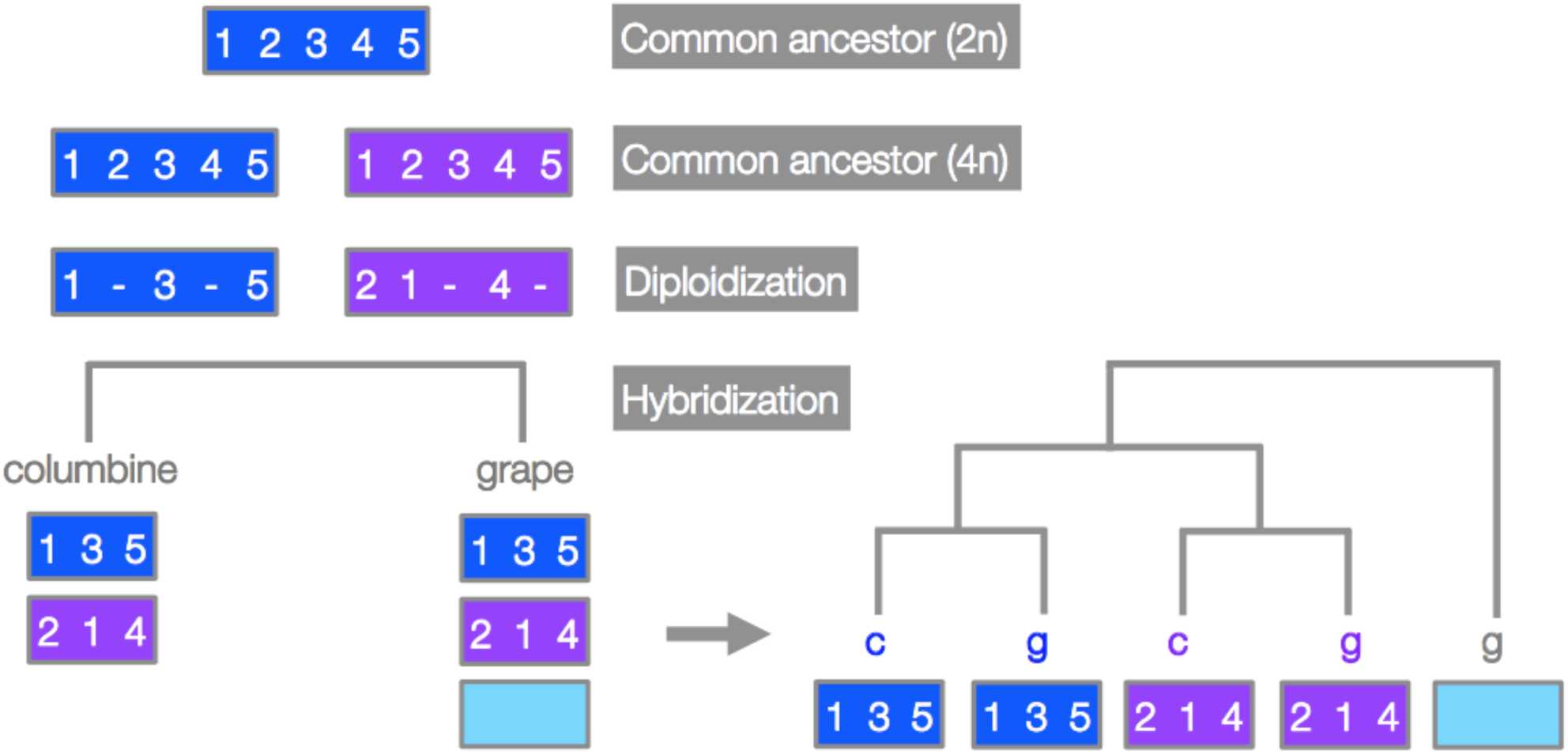
Gene order-based clustering expected under ancient tetraploidy common to all eudicots. Represented here by blue and purple rectangles, each member of the paralogous chromosome pair experiences intrachromosomal rearrangements as a part of the diploidization process. Deletions (denoted as “-”) will create the gene order ‘1, 3, 5’ on the blue chromosome while both deletions and an inversion will create the gene order ‘2, 1, 4’ on the purple chromosome. These differential paralogous gene orders will be inherited by both columbine and grape. If we compare the gene order on the homologous chromosomes of columbine and grape at this particular region, we should see “blue” chromosomes of columbine and grape forming one cluster while “purple” chromosomes of columbine and grape forming another cluster. Note that we show here only the “allohexaploidy” model, which predicts that the third grape paralog added via hybridization is an outgroup in this clustering analysis. See Fig. S11 for the expected gene orders under the “autohexaploidy” model.

We used two different approaches to detect this pattern. First, we clustered homologous segments based on gene-order similarity (Materials and Methods). The pairwise comparisons show that each member of columbine paralogs matches a different grape chromosome (Figs. S6-8). Reshuffling genes on grape chromosomes further indicates that this pattern of clustering is highly unlikely to be produced by chance (p=0-0.05). Second, we attempted to corroborate the clustering based on gene-order similarity by clustering homologous regions based on similarity in protein sequence (Materials and Methods). Because of the deep history of shared tetraploidy, only a small fraction of all the informative gene trees (0.016-0.044) show the “expected” pairings (Figs. S6-8), and it is thus not possible to infer history from individual trees. However, the order of the homologous genes that do show the expected pairwise clustering (based on sequence divergence) again recaptures the clustering pattern inferred from synteny alone (compare Figs. S7 and S9). Thus, the clustering pattern inferred from synteny is mirrored in clustering based on sequence divergence.

An eudicot-wide WGD is further supported by the observation that a chromosomal fusion, presumably experienced by the common tetraploid ancestor, is still detectable in the genomes of columbine and grape despite their separation of around 125 million years [37]. The first hint comes from the composition of the chromosomes: columbine chromosome 5 and grape chromosome 7 both appear to be fusions of the same ancestral chromosomes (Fig. 6). If these chromosomes were created by a single fusion event in the common tetraploid ancestor of eudicots, they should match each other with respect to gene order on each of the two homologous portions (“orange” and “green” portions in Fig. 7). This is what we see: columbine chromosome 5 and grape chromosome 7 cluster together with respect to their gene order on the “orange” portion (Fig. S7). For the “green” portion, columbine chromosome 5 matches grape chromosome 4 (Fig. S8), which used to be fused to grape chromosome 7 [38]. Additional support for shared ancestral fusion comes from the cacao (*Theobroma cacao*) genome [39]. The first chromosome of cacao does not only show a similar pattern of chromosomal ancestry [38,39], but also shares the gene order exclusively with the grape chromosomes 4 and 7 on the corresponding homologous portions (Fig. S10). In summary, the columbine fusion clusters with that of grape, which, in turn, clusters with that of cacao, strongly favoring a common origin of the fusion between “orange” and “green” ancestral chromosomes (Fig. 6).

**Fig. 6:**
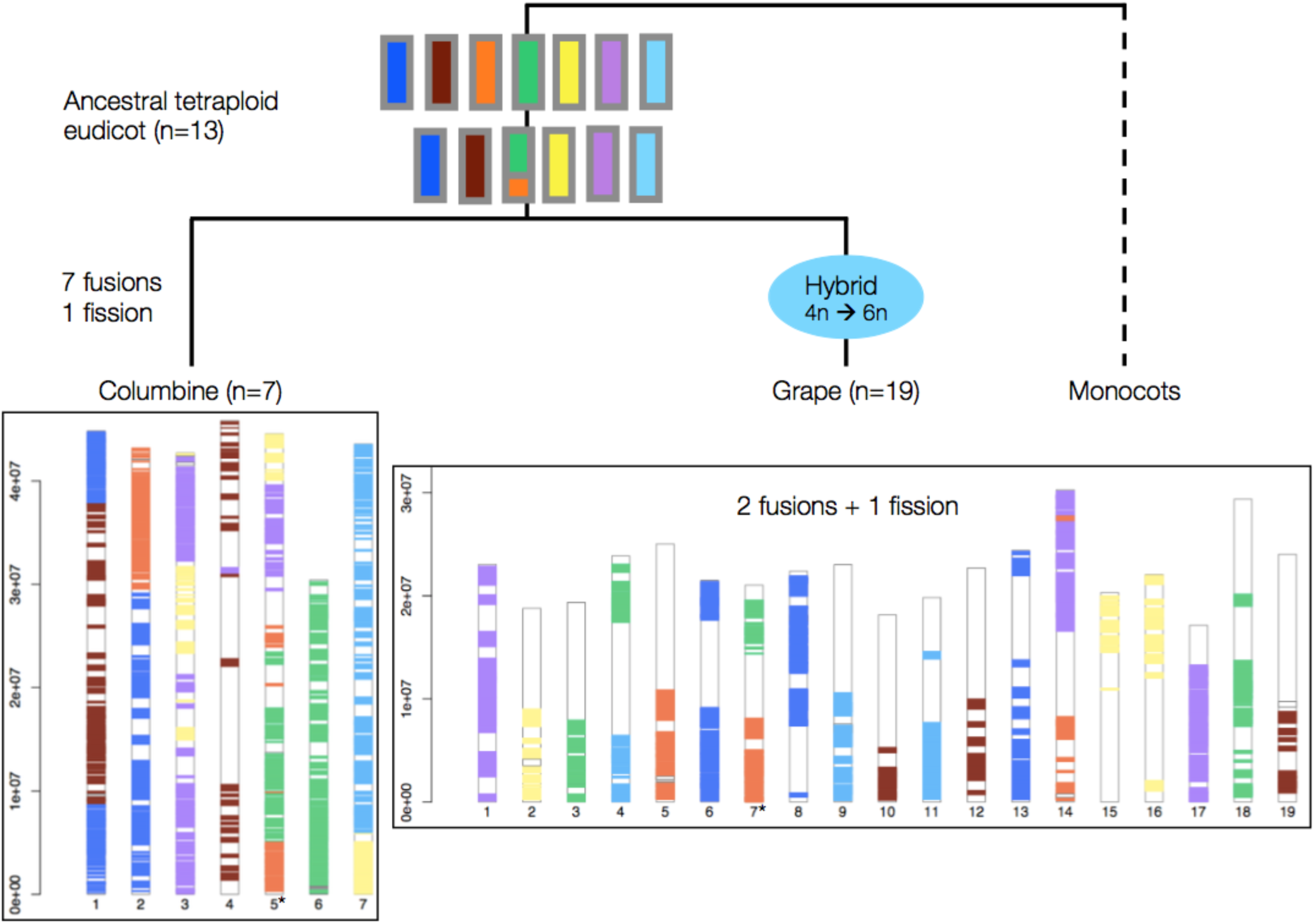
Tracing the genome reshuffling in columbine following tetraploidy. Grape chromosomes **(bottom right)** are colored by within-genome synteny. Seven distinct colors represent the haploid set of seven ancestral chromosomes before the eudicot-wide WGD. Each color is shared by three grape chromosomes reflecting the triplicate genome structure of core eudicots. The only exception is the “green” chromosome which is shared by four grape chromosomes due to a fission event [38]. Columbine chromosomes **(bottom left)** are colored by their synteny to grape chromosomes. Each color is generally shared by two chromosomes, reflecting columbine paleotetraploidy. As few as 7 fusions and a single fission are enough to explain the current structure of the columbine genome. Of these 7 fusions, 5 are between different chromosomes while 2 are between WGD-derived paralogous chromosomes. Columbine chromosomes 3 and 7 are examples of the latter (Figs. 1 and S4). Note that chromosome 5 of columbine and chromosome 7 of grape (*) both have the colors “orange” and “green” (cf. Fig. 7).

**Fig. 7:**
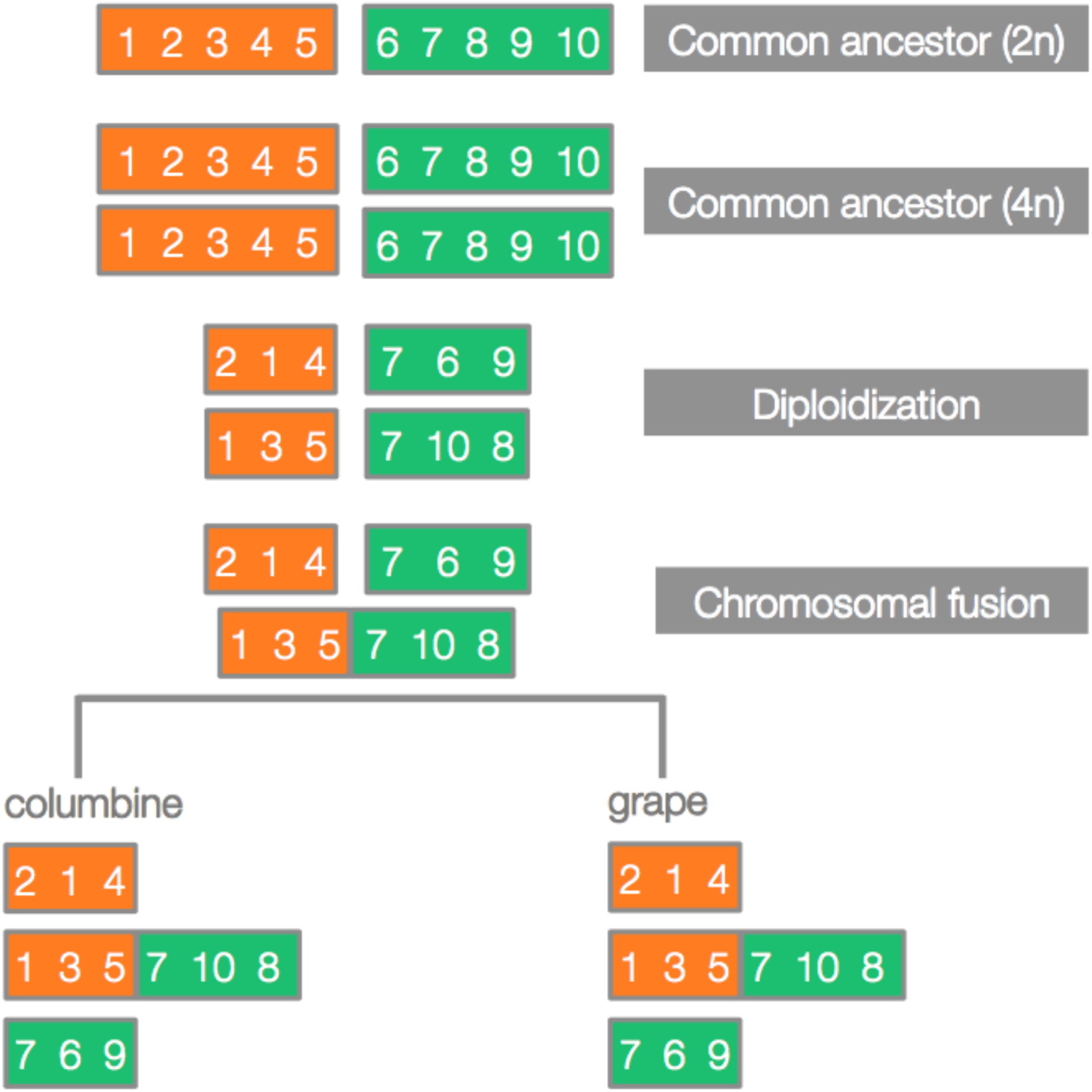
Schematic of predicted synteny patterns in the case of shared ancestral fusion. Two ancestral chromosomes (orange and green rectangles, with genes depicted as numbers) undergo WGD. Paralogous chromosome pairs diverge as a part of the diploidization process. A fusion joins one version of the “orange” chromosome (‘1, 3, 5’) with one version of the “green” chromosome (‘7, 10, 8’). If this took place in the common tetraploid ancestor of eudicots, the fused chromosomes in columbine and grape should also carry these versions on their “orange” and “green” portions. In the hypothetical example here, diploidization precedes the fusion event but may well happen afterwards with no effect on the predicted synteny patterns.

### The core eudicots have a hybrid origin

Our inference of shared tetraploidy between basal and core eudicots makes use of the signals presumably generated by diplodization (Figs. 5 and 7). However, hybridization of unreduced gametes from two divergent diploid genomes, “allotetraploidy”, would also lead to gene order-based clustering between two different pairs of grape and columbine chromosomes (Figs. S11-12). In this case, the alternative paralogous gene orders in the tetraploid ancestor reflect the gene orders on the progenitor chromosomes. Thus, the clustering pattern does not depend on whether the eudicot tetraploid genome evolved via “auto-” or “allopolyploidy”. The same is not true for the second part of the process leading to hexaploidy. In this case, autopolyploidy would lead to the duplication of one of the existing chromosomes, and only allohexaploidy would lead to one of the three paralogous grape chromosomes being an “outlier” with respect to the two grape-columbine pairing (Figs. S11-12) — which is what we see in our data (Figs. S6-8).

If our interpretation is correct and all core eudicots have a hybrid origin, the pattern of gene order-based clustering should be conserved. That is, we should be able to identify the hexaploidy-derived “outlier” chromosomes in other core eudicot genomes as well. To check this expectation, we again used the cacao genome, one of the most conserved genomes after grape [9,39]. Pairwise alignment between the homologous regions of columbine and cacao confirms our expectation: each member of columbine paralogs pairs up with a single cacao chromosome, leaving one of the cacao paralogs as an outlier (Figs. S13-14). Furthermore, as shown in Fig. 8 (see also Fig. S10), the cacao regions putatively derived from tetraploidy and hexaploidy, respectively, show a very clear one-to-one match to those in grape (detected in the grape-columbine comparison). As expected, the putatively orthologous pairs of cacao and grape regions show similar levels of synteny conservation with their paralogous counterparts, with the “outlier” regions being the most divergent [38]. Thus, although not independent evidence, the cacao genome supports a hybrid origin, and highlights the key role of the columbine genome in unravelling the history of the eudicot genome.

**Fig. 8:**
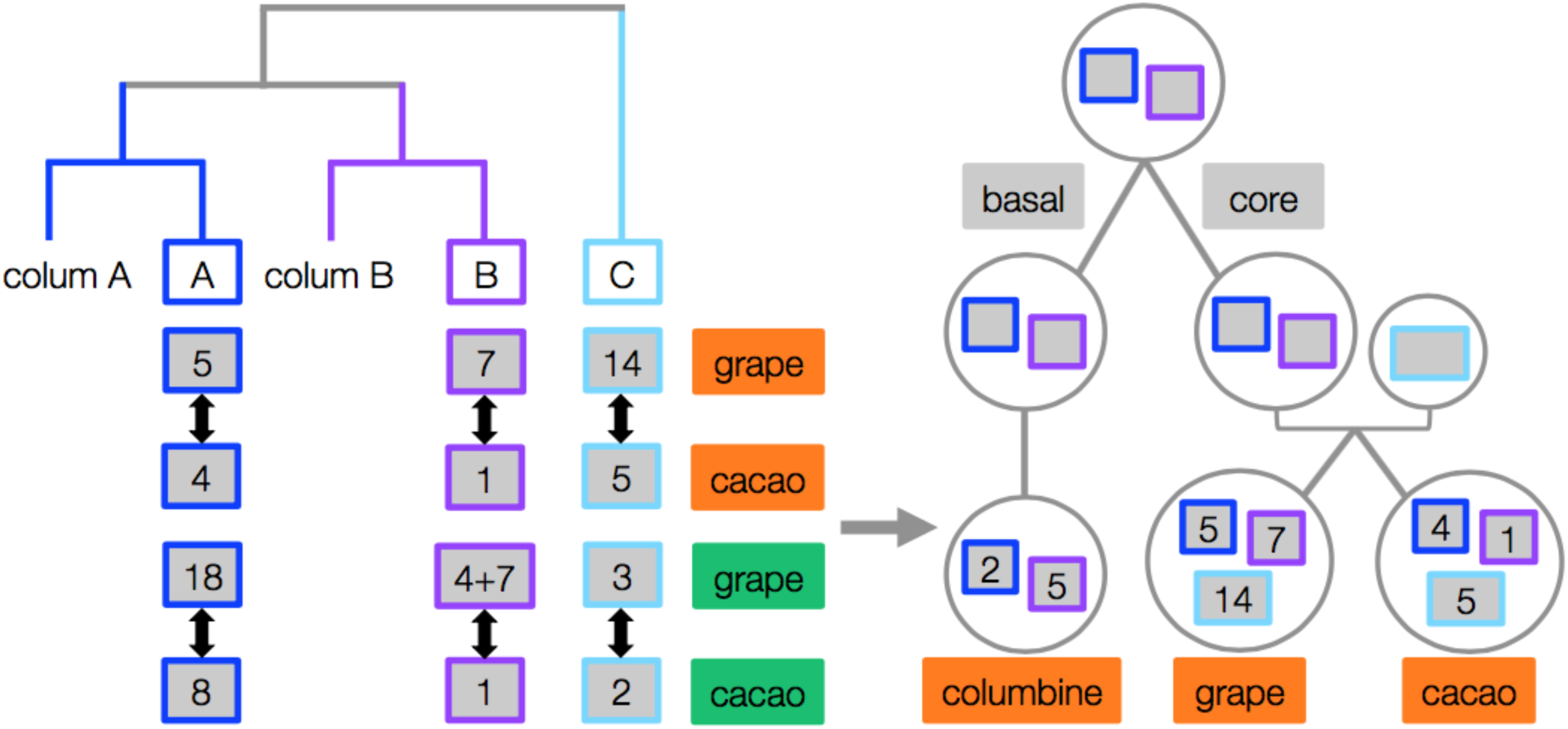
The shared history of chromosomes in columbine, grape and cacao. Gene order-based clustering results **(left panel)** are summarized here for the chromosomes harboring the “orange” and “green” homologous portions. The former corresponds to 5, 7, 14 in grape and 1, 4, 5 in cacao. The latter corresponds to 3, 4+7 (products of a fission), 18 in grape and 1, 2, 8 in cacao. In columbine, the “orange” portions are on chromosomes 2 and 5 while the “green” portions are on chromosomes 6 and 5, each pair of which being denoted as *colum A* and *colum B*, respectively. Both grape- and cacao-columbine pairing distinguish tetraploidy-derived regions (blue and purple rectangles) from hybridization-derived ones (light blue rectangles), defining the orthologous sets of regions across the three eudicot genomes **(right panel)**. The conservation of gene order exclusively between the putatively orthologous regions of grape and cacao (black arrows, Fig. S10) further strengthens our columbine-based inference of orthology.

### Current columbine chromosomes have mostly been generated via fusions

It is widely accepted that genome shuffling post-WGD has shaped the present-day karyotypes of all plant genomes [34]. Nevertheless, the extent of genome shuffling as a part of the “re-diploidization” process seems to vary widely: only 3 chromosomal rearrangements post-*gamma* are enough to explain the current structure of the grape genome (Fig. 6) while almost 150 chromosomal rearrangements were necessary for the sunflower genome to reach its current karyotype after several rounds of WGD [11]. To check where columbine falls in this spectrum, we identified chromosomal rearrangements likely to have happened after the tetraploidy shared by all eudicots: if the pre-WGD ancestral eudicot karyotype had a haploid number of 7 chromosomes [28], only seven columbine-specific fusions and a single fission are enough to explain the reduction in columbine chromosome number from n=13 to n=7 after the ancestral fusion event (Fig. 6). These rearrangements involve all the chromosomes in columbine apart from chromosomes 4 and 6, the former of which paradoxically shows the greatest erosion of synteny with grape chromosomes (Figs. 6 and S3). Given all the evidence suggesting a “decaying” nature of columbine chromosome 4 [19], we repeated the analysis of grape-columbine synteny detection with relaxed parameter settings. We did this by decreasing the minimum number of aligned gene pairs within a block (from 5 to 3) and increasing the maximum genic distance between matches (from 20 to 30). This allowed us to extend the synteny blocks towards more proximal regions (Fig. S15). Further zooming into the synteny relationship between grape chromosomes that are homologous to columbine chromosome 4 confirmed that there is no evidence of a fusion event (Fig. 9).

**Fig. 9:**
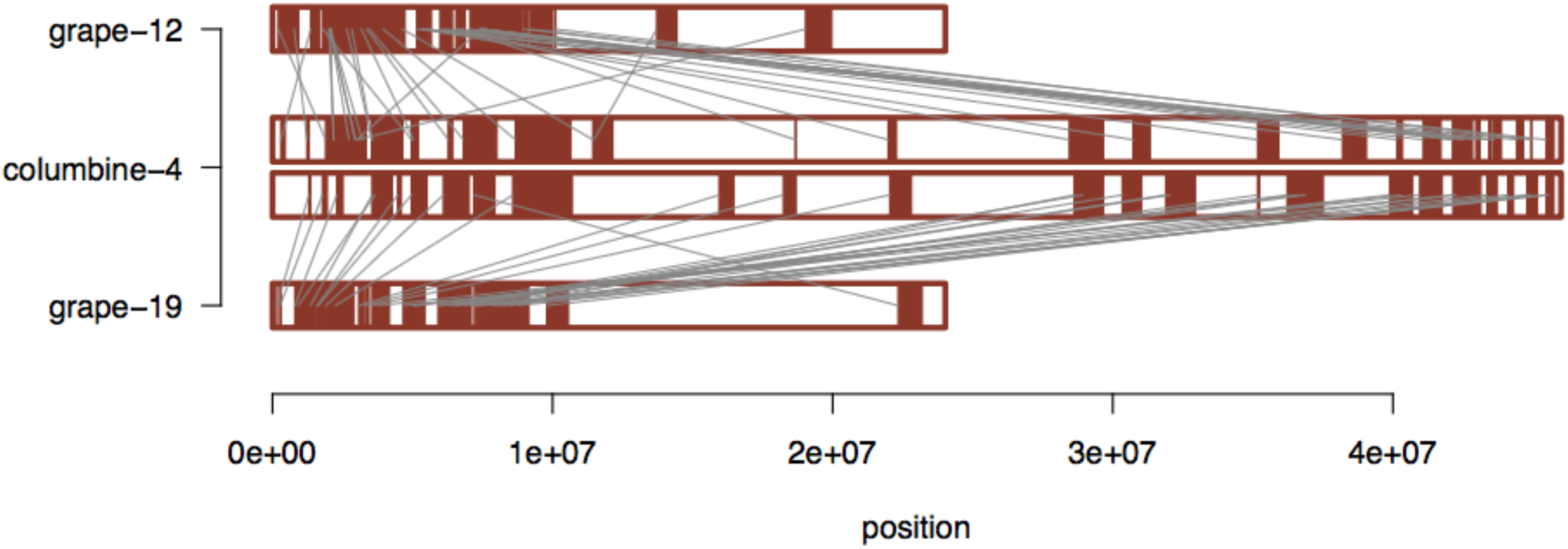
Synteny between columbine chromosome 4 and grape chromosomes 12 and 19. Much smaller grape chromosomes look like the compact versions of columbine chromosome 4. Note that this result is generated with the most relaxed parameter combination in Fig. S15, but holds true for a less relaxed combination of parameters as well (Fig. S16).

The lack of a fusion event on columbine chromosome 6 might explain the fact that it is the smallest chromosome of columbine (Fig. 6). However, chromosome 4 is comparable in size to the remaining chromosomes, all of which are products of ancient fusion events. The observations that chromosome 4 has a higher proportion of genes in tandem duplicates (0.37 versus genome-wide mean of 0.22) and a greater extent of intra-chromosomal synteny (indicative of segmental duplications) (Fig. S17) suggest that chromosome 4 has reached a comparable size partly due to numerous tandem and segmental duplications and partly due to an expansion of repetitive DNA [19]. These results reinforce the idea that chromosome 4 has followed a distinct evolutionary path from the rest of the genome.

Fusion-dominated genome shuffling [34] is not the only facet of diploidization [40]. Following WGDs, gene duplicates get lost and this happens in a non-random manner. Genes involved in connected molecular functions like kinases, transcription factors and ribosomal proteins are retained in pairs [41–45] potentially due to dosage-related constraints [46]: losing or duplicating some, but not all of these dosage-sensitive genes might upset the stoichiometric relationship between their protein products [47–49]. Consistent with this dosage balance hypothesis, columbine genes potentially retained post-WGD (1302 genes across 76 syntenic regions; Supplementary Data 1) are enriched for the GO categories “structural constituent of ribosome”, “transcription factor activity”, “translation” (p<0.001) and “protein tyrosine kinase activity” (p<0.01). Tandemly duplicated genes (n=6972), on the other hand, are depleted for the GO categories “structural constituent of ribosome”, and “translation” (p=10^-17), reflecting the role of dosage-related purifying selection.

## Discussion

All flowering plants are descended from a polyploid ancestor, and with only a few exceptions (e.g. *Amborella* from basal angiosperms [50]), all of them experienced at least one further round of WGD. Within-genome synteny (Fig. 1) shows that columbine is an example of the latter, confirming the conclusions from other studies [23,24,51]. Here we show that this columbine tetraploidy is a remnant of a WGD at the base of all eudicots and is thus far more ancient than previously thought [23,51]. Furthermore, we use this observation to argue that the hexaploidy shared by the core eudicots must have involved allopolyploidy, i.e., presumably hybridization between the ancestral tetraploid and a diploid species.

A eudicot-wide WGD has been suggested by several studies [26,35,36,52]. Our synteny-based approach solidifies these findings by demonstrating that the columbine and grape genomes have inherited the genome structure of a common tetraploid ancestor. That we can trace such an ancient tetraploidy is due to two facts. First, the genome structure of columbine is well-preserved and free from recent WGDs. Second, genomes provide much greater information than genes alone. Indeed, a recent study on another basal eudicot [51], the opium poppy, highlights these two facts. Having experienced a recent WGD (∼ 8 million years ago), the genome of the poppy is dominated by syntenic gene pairs of low divergence (Fig. 1C in [51]), although it also carries highly diverged paralogs whose Ks values nicely overlap with our estimates for columbine and grape, consistent with an eudicot-wide WGD (compare Fig. S13D in [51] to Fig. 4). However, the strength of the signal from recent polyploidization largely obscures the much weaker signal of ancient polyploidization (interpreted as segmental duplications by Guo et al. [51]). In fact, although overlooked by the authors, the intergenomic synteny between columbine and poppy provides a clear signature of an eudicot-wide WGD (Fig. S9D in [51]). Differing from columbines with only one additional genome duplication, the poppy genome aligns to the columbine genome in a 4:2 manner, with 4 paralogous regions of poppy syntenic to 2 paralogous regions of columbine derived from the ancient shared tetraploidy.

Our approach also helps us shed light on the nature of the *gamma* hexaploidy found in all core eudicots [9,28–32, and Supplementary Note 5 in 33]. WGDs have often been discussed as if they were “events”, ignoring the process by which they originated. We show here that core eudicot hexaploidy is the result of two processes: an ancient tetraploidization shared by all eudicots, followed by allopolyploidization leading to the core eudicots. In other words, all core eudicots have a hybrid origin. An allohexaploid origin has indeed been previously suggested by Murat et al. [9], who identified the three subgenomes of grape using differential patterns of gene loss on “dominant” versus “sensitive” subgenomes. Their classification assumes that the most recently added set of paralogous chromosomes will be “dominant”, because they have spent a shorter amount of time in the polyploid genome and thus experienced fewer gene losses. Contrary to this, our results suggest that the most recently added grape chromosomes (chromosomes 3, 8, 9 and 14) largely corresponds to the “sensitive” grape chromosomes identified by Murat et al. [9]. Instead, we argue that the extensive gene loss in the most recently added subgenome reflects its divergence from the other two subgenomes at the time of hexaploid formation, perhaps similar to the situation in the allotetraploid *Arabidopsis suecica*, which is a hybrid between the more ancestral-like (n=8) genome of *A. arenosa*, and the heavily reduced (n=5) genome of *A. thalian*a [53]. Another example is hexaploid wheat, which is a hybrid between tetraploid emmer wheat and wild diploid grass, *Aegilops tauschii* [54 and references therein].

## Conclusions

Our findings reveal the hybrid structure of core eudicot genomes and will hopefully help us understand what hybridization has meant for core eudicots — a group which comprises more than 70% of all living flowering plants [55]. What are the hybridization-coupled changes that has led to the current patterns of gene expression, methylation, transposable element density/distribution? All these questions call for additional genomes from basal eudicots which — as this study illustrates — have great values as outgroup to the core eudicots. More data will also allow the development of sophisticated analysis methods based on explicit models of the evolution of gene order, which our results suggest is a very powerful source of information about the past.

## Materials and Methods

### Synteny detection

We performed all genes (CDS)-against-all genes (CDS) BLAST for the latest version of *Aquilegia coerulea* reference genome (v3.1) using SynMap tool [29] in the online CoGe portal [56]. We also looked at the synteny within *Vitis vinifera* (v12) and between *A. coerulea* and *V. vinifera* using default parameter combinations in DAGChainer. We filtered the raw output files for both within grape and columbine-to-grape synteny. For the former, we only kept the blocks that are syntenic between the polyploidy-derived paralogous chromosomes of grape as identified by Jaillon et al. [28] (Supplementary Data 3). For the latter, we required that a given columbine chromosome is overall syntenic to all the three paralogous chromosomes of grape (Supplementary Data 2). So, for a given pair of columbine and grape chromosomes, we only kept the blocks if the columbine chromosome also matches to the other members of paralogous grape chromosomes.

The raw output files can be regenerated at the CoGe portal [56] using the id numbers provided below for each species (Availability of data and material) and changing the default parameter combination in DAGChainer (D:A=20:5) when needed. D and A specify the maximum genic distance between two matches and the minimum number of aligned gene pairs, respectively, to form a collinear syntenic block.

### Estimating the divergence between synteny block pairs

We used Ks (the number of synonymous substitutions per synonymous site) values provided for each homolog gene pair by the CoGe portal [56]. We estimated the median Ks of homologous genes in a synteny block after filtering gene pairs with Ks>10 due to saturation effect [57]. Both values are provided in Supplementary Data 1-3 for within columbine, columbine-to-grape and within grape synteny, respectively.

### Quantifying gene order similarity

We “reconstructed” a given set of columbine and grape chromosomes at their homologous regions (color-coded in Fig. 6). We seeded this reconstruction by focusing on at least three consecutive genes aligning between a pair of columbine and grape chromosomes (D:A=0:3). We particularly chose three genes since it is the most stringent value we could use to detect homologous synteny blocks; we detected almost nothing when we required 4 consecutive genes (D:A=0:4). This stringent criteria aim to minimize the effect of gene movement on the homology between columbine and grape chromosomes. Once we had the list of genes, we then looked for their paralogous counterparts on the remaining columbine and grape chromosomes using intragenomic gene-to-gene blast (D:A=0:1). Having chromosomes represented by syntenic gene sets and reminiscent of these sets (Fig. S5 and Supplementary Data 4), we assigned a unique word to each synteny block and the genes forming the block to be able to use the text alignment provided by the R package *align_local* [58]. We then quantified the gene (“word”) similarity as such: for an initial N number of words on a columbine chromosome (N=window size), we did a pairwise alignment between these N words and all the words a grape chromosome (match=4, gap=-1). We repeated the same analysis with the inverted order of N words and picked the maximum alignment score. We repeated these steps by sliding the window by one word and keeping the N constant to get a distribution of scores as in Figs. S6-8. We used different N values ranging from 4 to 15. Note that we excluded columbine chromosomes 3, 4 and 7 from this analysis since they all have a complex history of lineage-specific chromosomal reshuffling events (Figs. 1, 6, S4 and S17).

We applied the same stringent criteria (D:A=0:3) to detect the homologous regions between grape and cacao (*Theobroma cacao*, v1). The same criteria led to very few homologous regions between columbine and cacao. So, we relaxed the parameters for the synteny detection between these two genomes (D:A=0:2) and quantified the gene order similarity with greater window sizes (N=20, 30, 35, 40 and 50). Note that we focused on the triplicated regions distributed across 3 different cacao chromosomes (Figs. 8, S13-14), which are rather unaffected by lineage-specific shuffling [38].

### Statistical testing of gene order similarity

Given the gene order similarity between the two different pairs of columbine and grape chromosomes harboring homologous regions, we performed permutation tests to estimate the probability of observing such a clustering just by chance. To do so, we first combined all the grape genes and sampled the same number of genes (“words”) as we observe to reconstruct each of the paralogous grape chromosome. We repeated the quantification step as above to get a permuted distribution of alignment score between a pair of columbine and grape chromosomes. We used Wilcoxon rank sum test (W-statistic) to quantify the shift in the distribution of alignment scores between one of the members of columbine paralogous chromosomes and its best grape hit when combined with the alignment scores between the same columbine chromosome and other grape chromosomes. We repeated the same analysis for the other member of columbine paralogous chromosomes as well. Having these *observed* W-statistics, we counted the number of cases (out of 100) where the permuted distributions generate W-statistics as high as or higher than the observed ones. We ran permutation tests for the columbine-cacao pairing as well (Figs. S13-14).

### Building gene trees

We built upgma trees for the homologous genes (Supplementary Data 5) distributed across a given set of columbine and grape chromosomes (color-coded in Fig. 6). We first detected homologous genes aligning between a pair of columbine and grape chromosomes (D:A=0:1). We then searched for their paralogous counterparts using intragenomic blasts (D:A=0:1). For protein alignment, we required at least five homologous genes, each from a single chromosome in the given set and ran ClustalW2 (v2.1) with the options *-TREE -KIMURA -CLUSTERING=UPGMA -OUTPUTTREE=dist* [59]. Of all the trees generated by ClustalW2 (informative trees), we only focused on the ones that support the synteny-based pairings (Figs. S6-8), which are detected by the *subtrees* function in R package *ape* [60,61]. Once we had the sets of homologous genes from this subset of trees, we assigned a unique word to the each set and quantified the gene order similarity between pairs of columbine and grape chromosomes as mentioned above.

For protein sequences, we used the annotations provided by JGI and Ensembl for columbine [19] and grape [62], respectively. Note that CoGe [56] outputs grape genes with the “PAC” tag while they are tagged with “VIT” in the Ensembl database. To match these different ids, we used two intermediary files. The first one is a gff file provided by CoGe (available at https://genomevolution.org/coge/GenomeInfo.pl?gid=19990). The second one is a conversion file provided by the Grape Genome Database [63] that lists the correspondence between different gene ids (can be downloaded from http://genomes.cribi.unipd.it/DATA). These two files contain the common tag “GSVIVT” which bridges the “PAC” and “VIT” tags.

### GO enrichment analysis

We used gene annotations provided by JGI [19] to test the null hypothesis that the property for a gene to be retained post-WGD and to belong to a given GO category are independent. We created a 2×2 contingency table as shown below and applied Fisher’s exact test for each GO category independently. We repeated the same analysis for tandem gene duplicates as identified by SynMap [29,56]; this time testing the null hypothesis that the property for a gene to be tandemly duplicated and to belong to a given GO category are independent. We excluded genes on scaffolds and reported enriched/depleted categories if they remain significant (p <0.05) after multiple test correction (fdr).

**Table 1:** 2×2 contingency table obtained by classifying genes into 2 categorical variables. The letters denote the number of genes for a given category (e.g. “a” denotes the number of retained genes annotated with the tested GO category).

|  | GO | not-GO | SUM |
| --- | --- | --- | --- |
| retained | a | b | a+b* |
| not-retained | c | d | c+d |
| SUM | a+c | b+d | N=total number of genes<br>=29550 (across 7 chromosomes) |

\*equal to 1302 and 6972 for candidate WGD-derived paralogs and tandem gene duplicates, respectively.

## List of abbreviations

WGD: whole genome duplication;
Ks: the number of synonymous substitutions per synonymous site;
GO: Gene Ontology.

## Declarations

### Availability of data and material

The columbine, grape and cacao genomes are available at the CoGE portal for the synteny analyses with the id numbers 28620, 19990 and 25287, respectively [56].

### Authors’ contributions

G.A. performed all analyses. G.A. and M.N. wrote the manuscript.

#### Acknowledgements

We thank Robin Burns and Claus Vogl for their comments on the manuscript; Daniel Gómez Sánchez and Benjamin Jaegle for the fruitful discussions.

