## Supplemental Figures for "The *Aquilegia* genome reveals a hybrid origin of core eudicots"

### Supplementary Figures

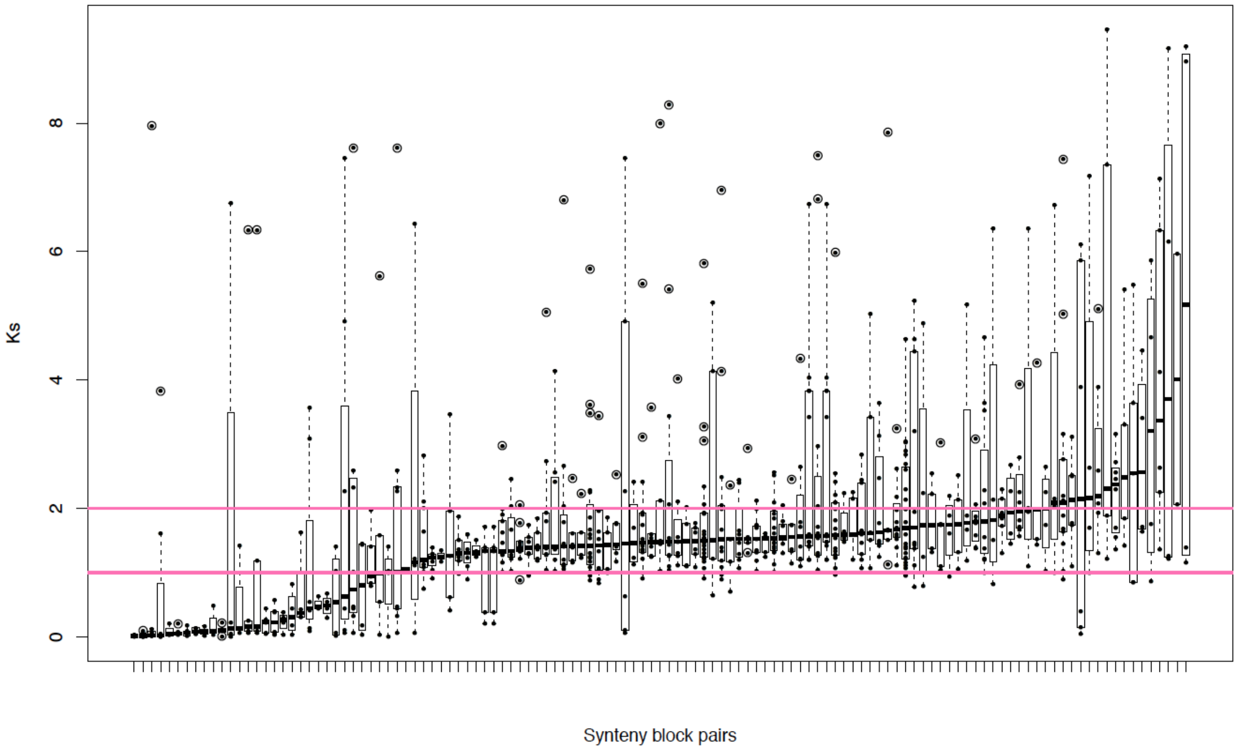


**Fig. S1: Distribution of Ks between duplicated genes located in synteny block pairs.** The block pairs (121 in total) are sorted with respect to the median Ks of gene duplicates. Pink lines refine block pairs (76 in total) with median Ks falling between 1 and 2 (Supplementary Data 1). Encircled points denote outliers.

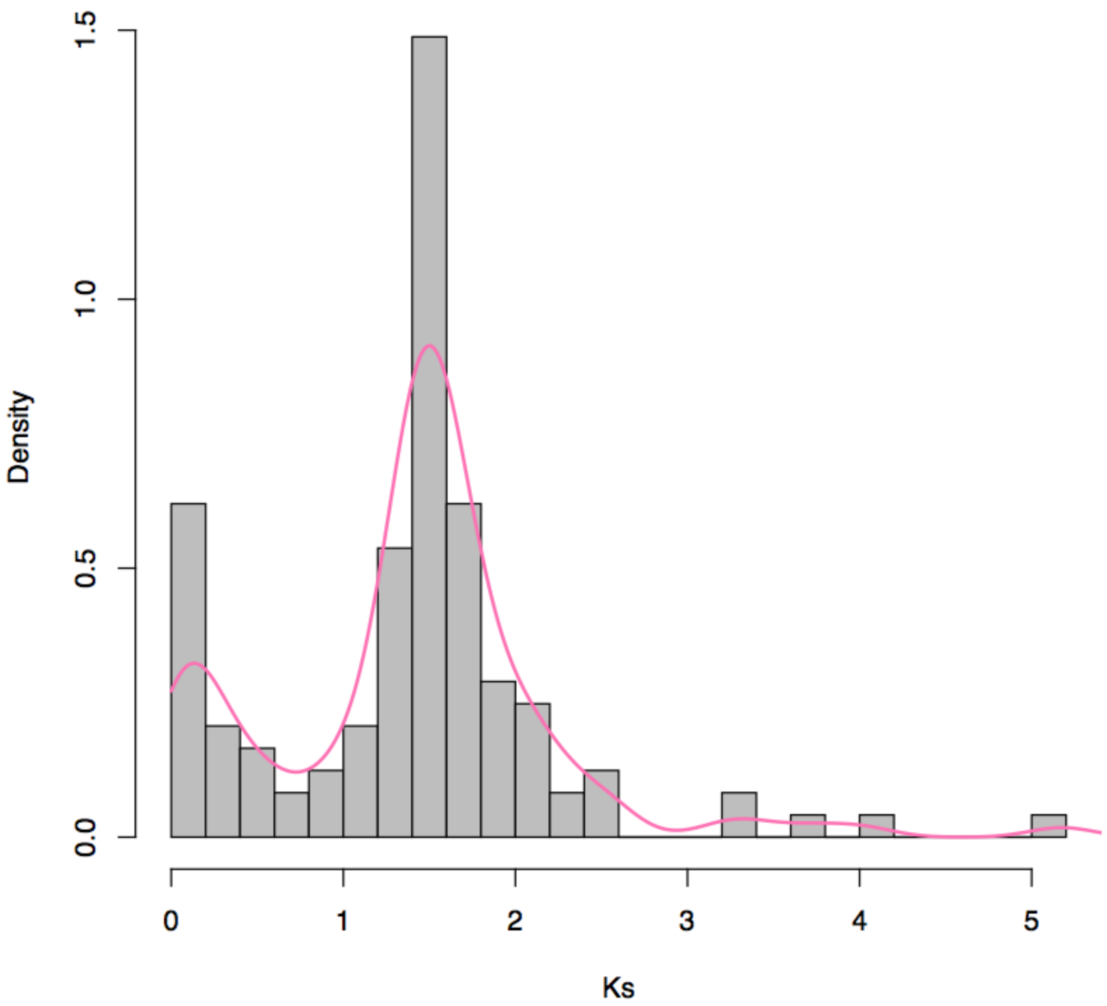


**Fig. S2: Age distribution of synteny blocks.** Measured as median Ks of all duplicate gene pairs in a syntenic block, the “age” of blocks clusters around Ks=1-2, producing the peak of gene birth expected from an ancient polyploidy. The initial peak (Ks < 0.2), on the other hand, most probably reflects recent small-scale duplications. Consistent with this, 6 out of 15 blocks under the initial peak are on the same chromosome (Supplementary Data 1), suggestive of recent intrachromosomal segmental duplications. Note that one of these “young” blocks is detected on *Aquilegia* chromosome 3, which has been generated via the fusion of WGD-derived paralog chromosomes (Fig. 1). So, gene conversion between paralogous regions, rather than a recent segmental duplication, might also result in a low Ks value.

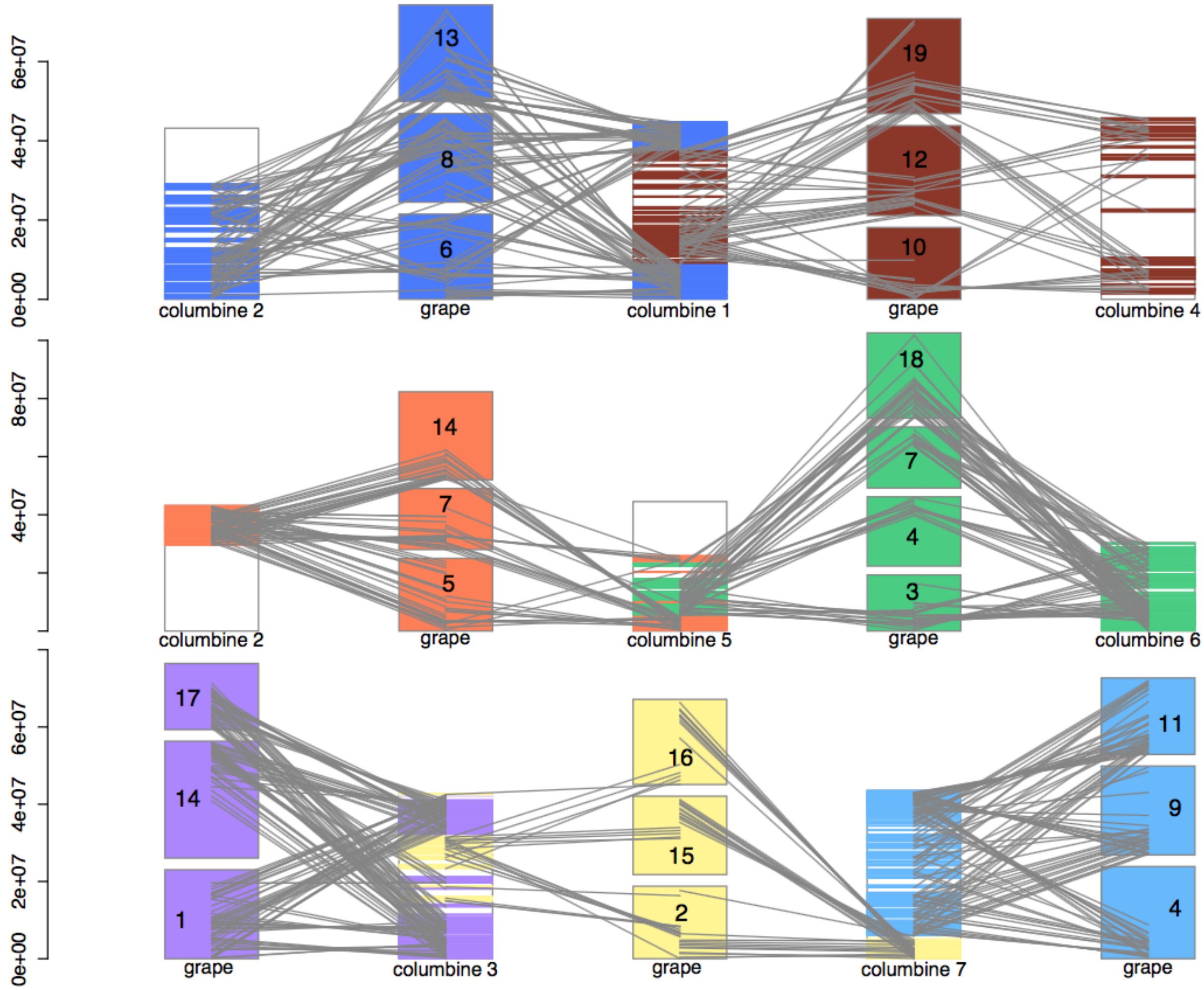


**Fig. S3: Synteny between columbine and grape chromosomes.** Grape is annotated with a haploid chromosome number n=19 [1]. Each set of WGD-derived paralog chromosomes in grape and the columbine region that is syntenic to it are colored uniquely (Supplementary Data 2). Grey lines further parse the synteny and reveal that two columbine chromosomes share homology with three grape chromosomes (first two rows), arguing against paleohexaploidy in columbine. This 2:3 ratio is less obvious for columbine chromosomes 3 and 7 (last row), each of which seem to have been created via two fusion events: a fusion between WGD-derived paralog chromosomes (Figs. 1 and S4) and another fusion with the "yellow" chromosome. Note that the end of columbine chromosome 5 is homologous to "purple" and "yellow" grape chromosomes (Fig. 6) but not shown here for simplicity.

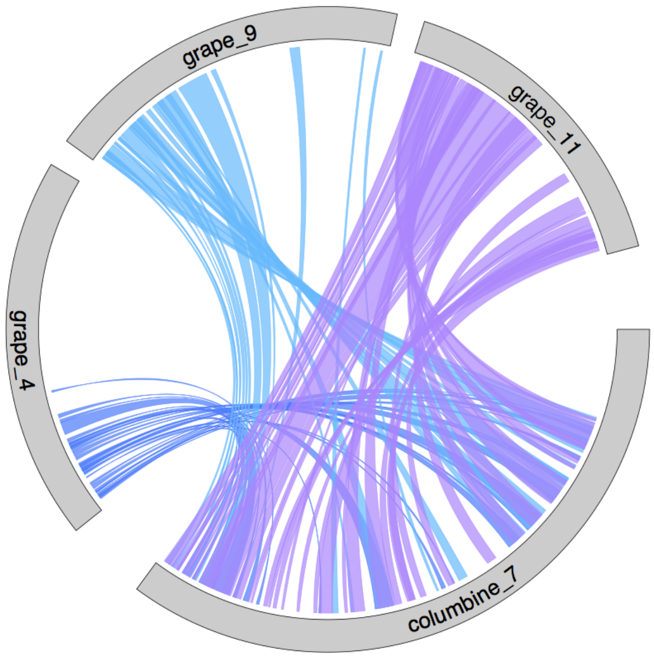


**Fig. S4: Synteny between the homologous regions of columbine and grape in the case of columbine-specific fusion.** The circos plot zooms into the synteny between columbine chromosome 7 and paralogous grape chromosomes 4, 9 and 11 (Supplementary Data 2) to show that both ends of columbine chromosome 7 align to the same region of a given grape chromosome as expected from the fusion of paralogous chromosomes.

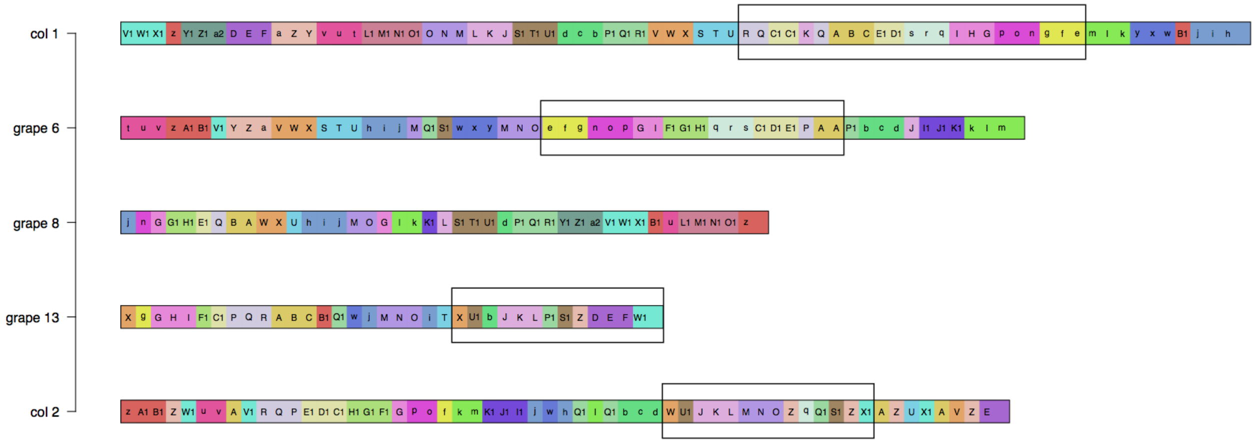

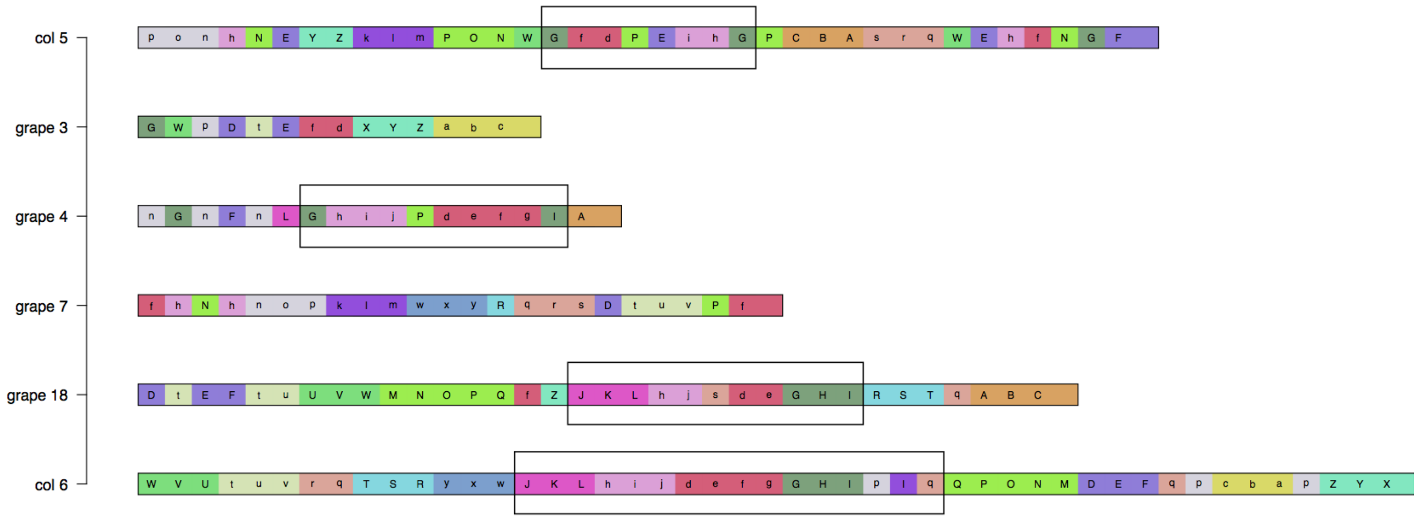


**Fig. S5: Reconstruction of chromosomes from genes consecutively matching between columbine and grape.** A synteny block, defined by a minimum number of three consecutive genes matching between a pair of columbine and grape chromosomes harboring homologous regions, is both color- and letter-coded, which, in turn, defines the color and letter codes of the genes it harbors. Results are presented here for columbine chromosomes 1 and 2 **(top)**, and 5 and 6 **(bottom)** and grape chromosomes 6, 8, 13 and 3, 4, 7, 18, respectively. Black rectangles frame the regions where a pair of columbine and grape chromosomes uniquely shares the gene order. These pairs include columbine chromosome 1 and grape chromosome 6, and columbine chromosome 2 and grape chromosome 13 (top); columbine chromosome 5 and grape chromosome 4, and columbine chromosome 6 and grape chromosome 18 (bottom).

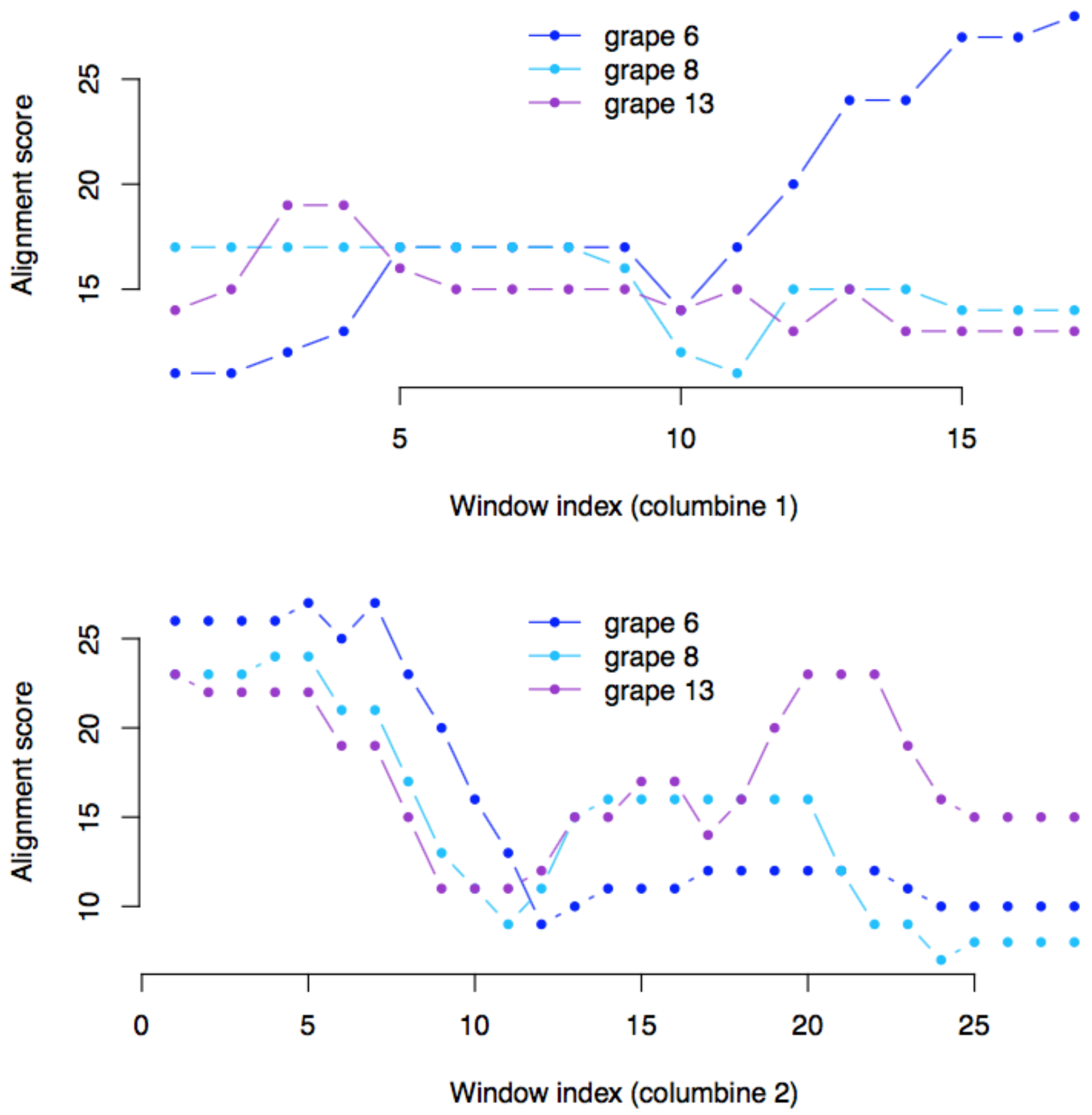


**Fig. S6: Examples of gene order similarity between the homologous regions of columbine and grape.** For successive windows of genes within a given columbine chromosomal region, the best alignment score with respect to each of the three grape chromosomes harboring homologous regions, is given. For example, columbine chromosomes 1 and 2 share a paralogous region homologous to grape chromosomes 6, 8, and 13 (Figs. 6 and S3). The chromosome 1 region **(top panel)** appears to be most closely related to grape chromosome 6, whereas its paralogous counterpart on chromosome 2 **(bottom panel)** appears to be most closely related to grape chromosome 13. See Fig. S5 for the correspondence between gene orders and scores, which peak towards the end of each columbine region. Note that the results presented here are shown for a window size of 12 genes but remain significant for all the window sizes tested (p=0-0.05).

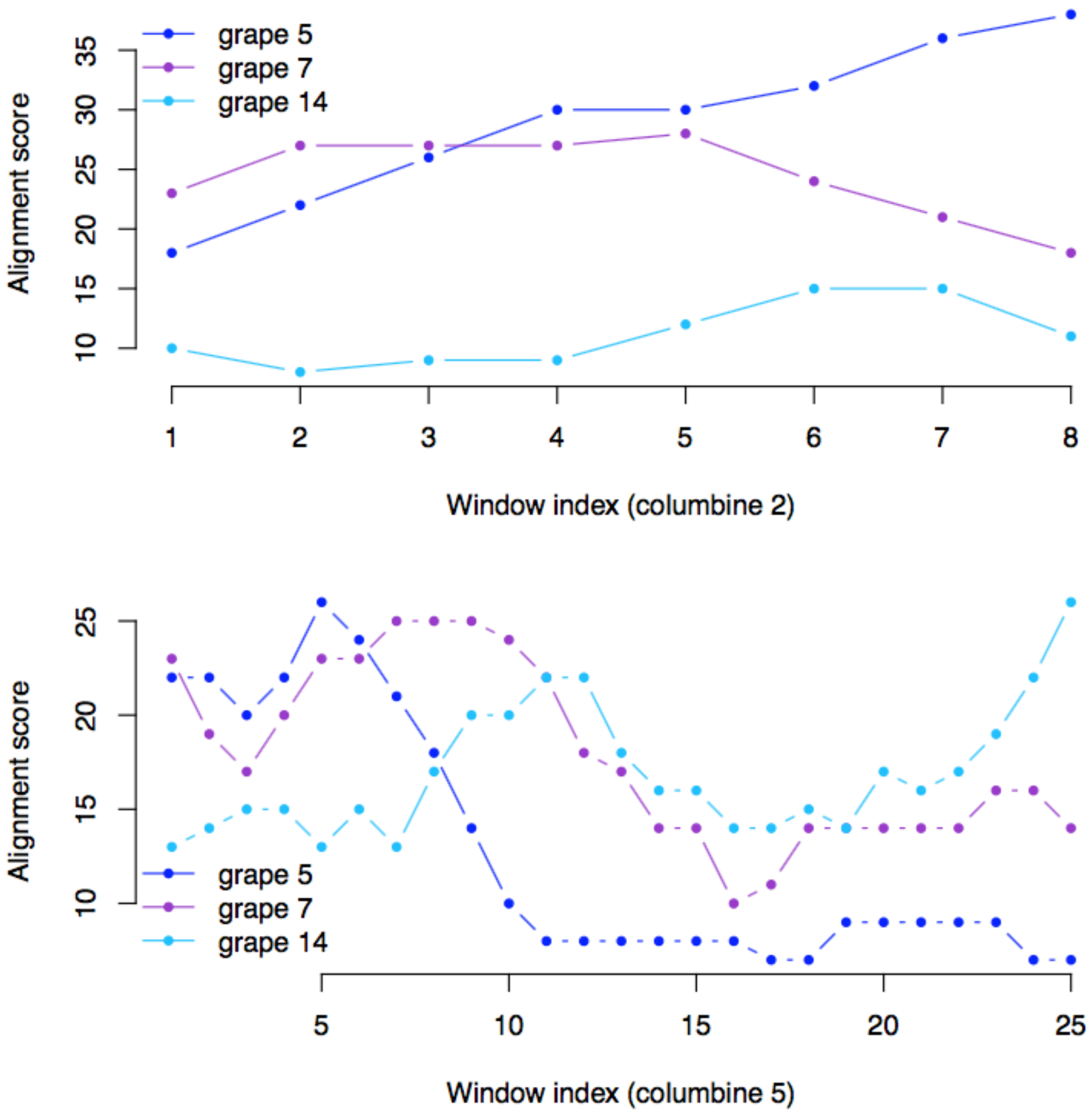


**Fig. S7: Examples of gene order similarity between the homologous regions of columbine and grape.** For successive windows of genes within a given columbine chromosomal region, the best alignment score with respect to each of the three grape chromosomes harboring homologous regions, is given. For example, columbine chromosomes 2 and 5 share a paralogous region homologous to grape chromosomes 5, 7, and 14 (Figs. 6 and S3). The chromosome 2 region **(top panel)** appears to be most closely related to grape chromosome 5, whereas its paralogous counterpart on chromosome 5 **(bottom panel)** appears to be most closely related to grape chromosome 7. Note the increased similarity between the chromosome 5 end region and chromosome 14 in columbine and grape, respectively, which potentially reflects the homology between the "purple" portions rather than the "orange" portions of focus here

(Fig. 6). Note also that the results presented here are shown for a window size of 10 genes but remain significant for all the window sizes tested (p=0-0.02).

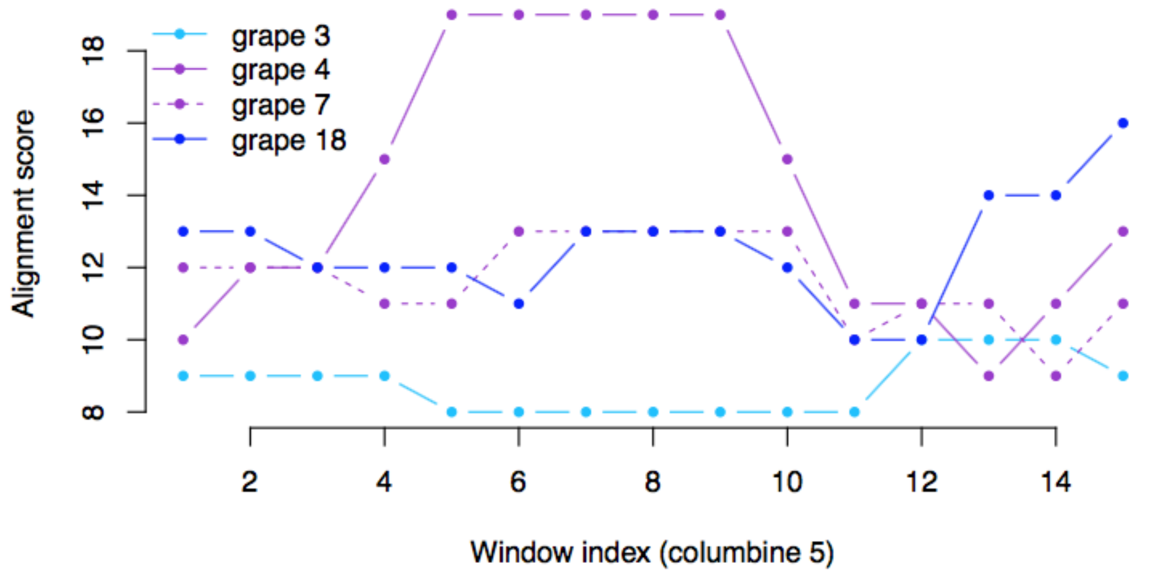


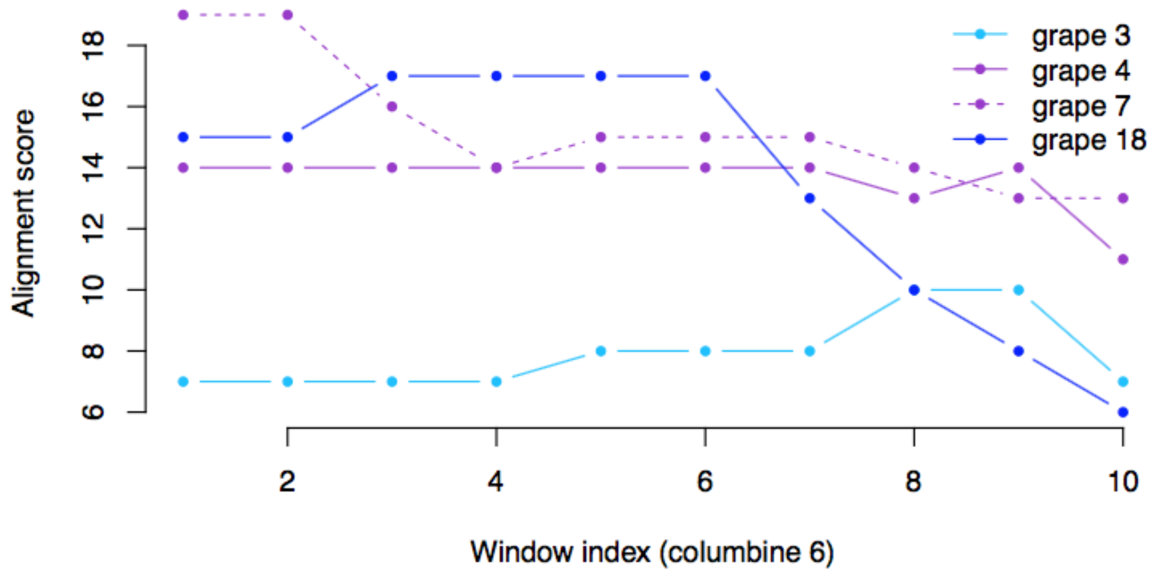


**Fig. S8: Examples of gene order similarity between the homologous regions of columbine and grape.** For successive windows of genes within a given columbine chromosomal region, the best alignment score with respect to each of the four grape chromosomes harboring homologous regions, is given. For example, columbine chromosomes 5 and 6 share a paralogous region homologous to grape chromosomes 3, 4, 7, and 18 (Figs. 6 and S3). The chromosome 5 region **(top panel)** appears to be most closely related to grape chromosome 4, whereas its paralogous counterpart on chromosome 6 **(bottom panel)** appears to be most closely related to grape chromosome 18. Note that the results presented here are shown for a window size of 10 genes but remain significant for all the window sizes tested (p=0-0.03).

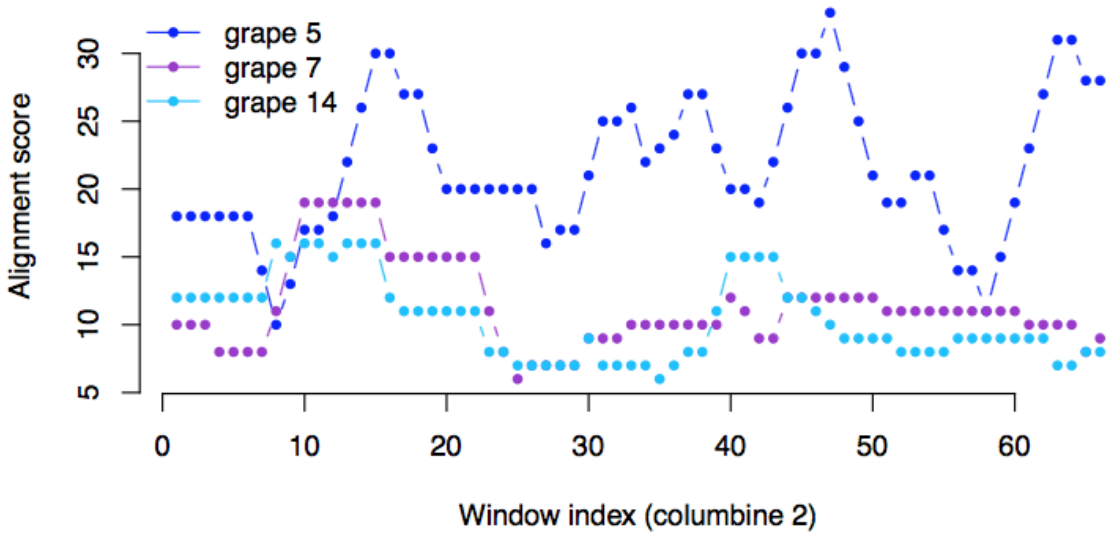


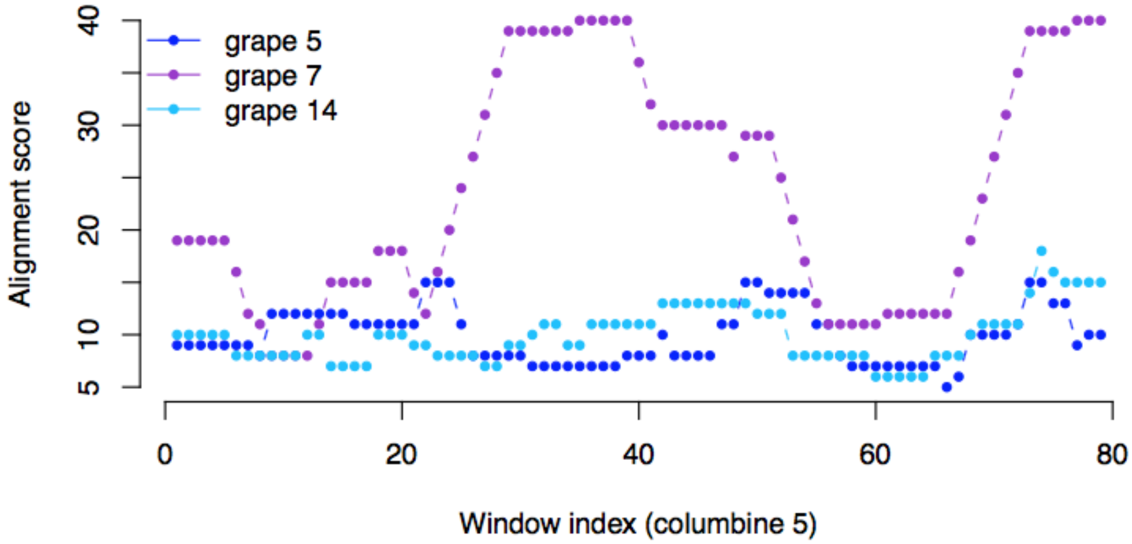


**Fig. S9: Examples of similarity in the order of homologous genes supporting the expected pairwise clustering.** The results shown here are for the same set of columbine and grape chromosomes and window size as in Fig. S7. This time, the pairwise similarity is based on the order of homologous genes that support the synteny-based pairing (Fig. S7). Note that the correspondence between synteny- and sequence-based pairings holds for the chromosome sets in Figs. S6 and S8. For the latter, we capture the signal from grape chromosome 7 instead of grape chromosome 4.

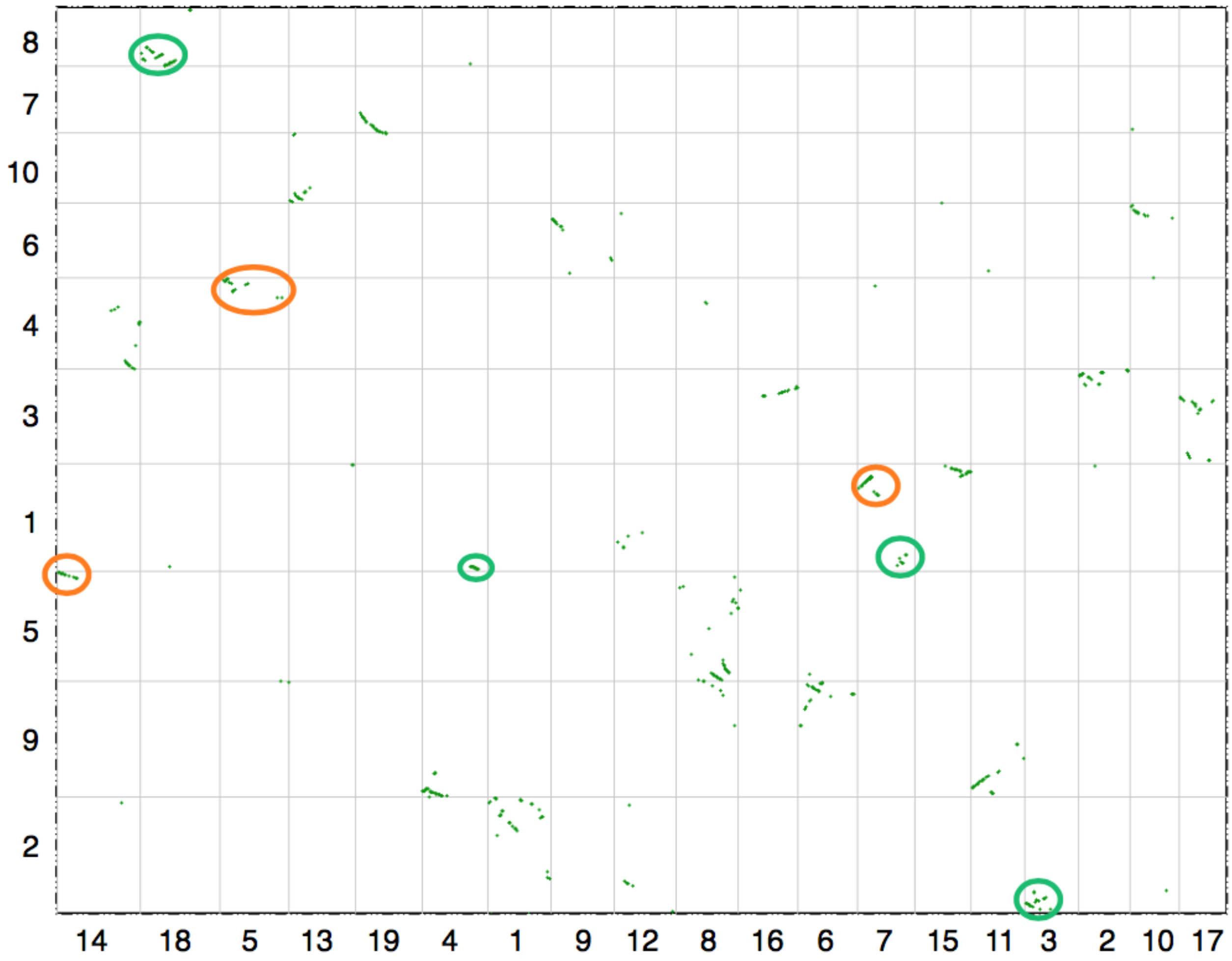


**Fig. S10: Synteny between cacao (y-axis) and grape (x-axis) chromosomes.** Shown here as a dot plot, each synteny block is required to harbor at least three consecutive gene pairs. The overall one-to-one match between the homologous regions of cacao and grape is highlighted for the "orange" and "green" portions of grape (Fig. 6). Note that cacao chromosome 1 shares the gene order with grape chromosomes 4 and 7 on the "green" portion, the former of which used to be fused to the latter [2].

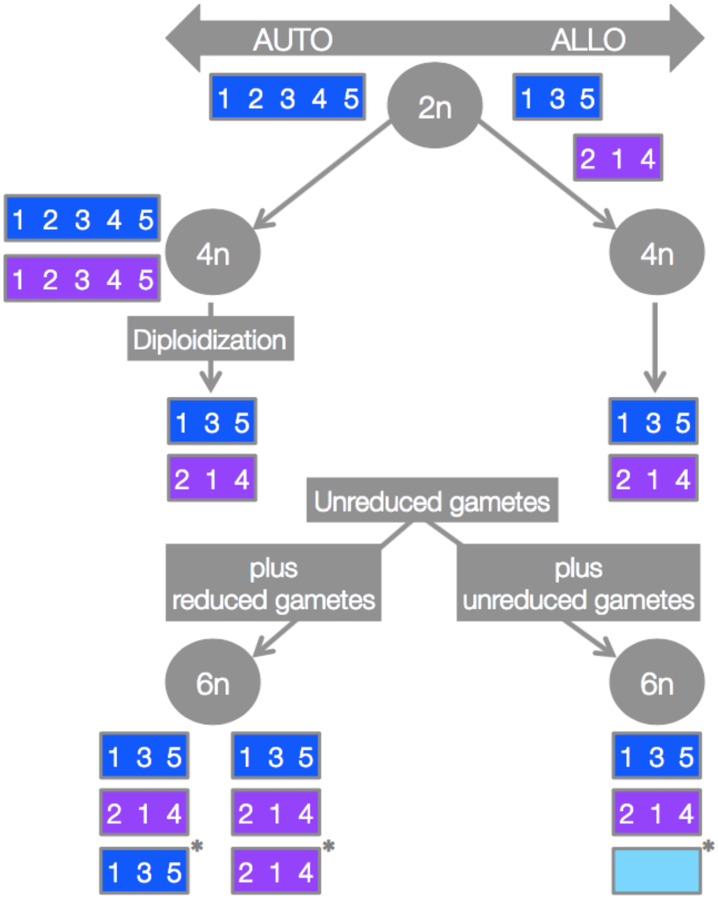


**Fig. S11: Expected gene orders on the three paralogous grape chromosomes under auto- versus allopolyploidy.** Starting from the common diploid ancestors of eudicots (2n), the production of unreduced gametes provides the substrate for polyploid formation. For each polyploidy level, only reduced gametes are shown with rectangles and numbers representing chromosomes and genes, respectively. Gametes with same gene order fuse under "autopolyploidy" **(left)** while gametes with different gene orders fuse under "allopolyploidy" **(right)**. Either mode can give rise to the alternative paralogous gene orders in the tetraploid genomes (4n). The mode of polyploidy, however, matters when it comes to hexaploidy. The third and most recently added chromosome (*) should share the gene order with one of the two other paralogous chromosomes under "autopolyploidy". If the third set of chromosomes is contributed by a different genome, as expected under "allopolyploidy", the most recently added chromosome should fall as an outlier with respect to its gene order (light blue rectangle). Note that, "autohexaploidy" involves the fusion of unreduced (4n) and reduced (2n) gametes while "allohexaploidy" involves the fusion of unreduced gametes, one from tetraploid progenitors (4n) and the other from genetically different diploid progenitors (2n).

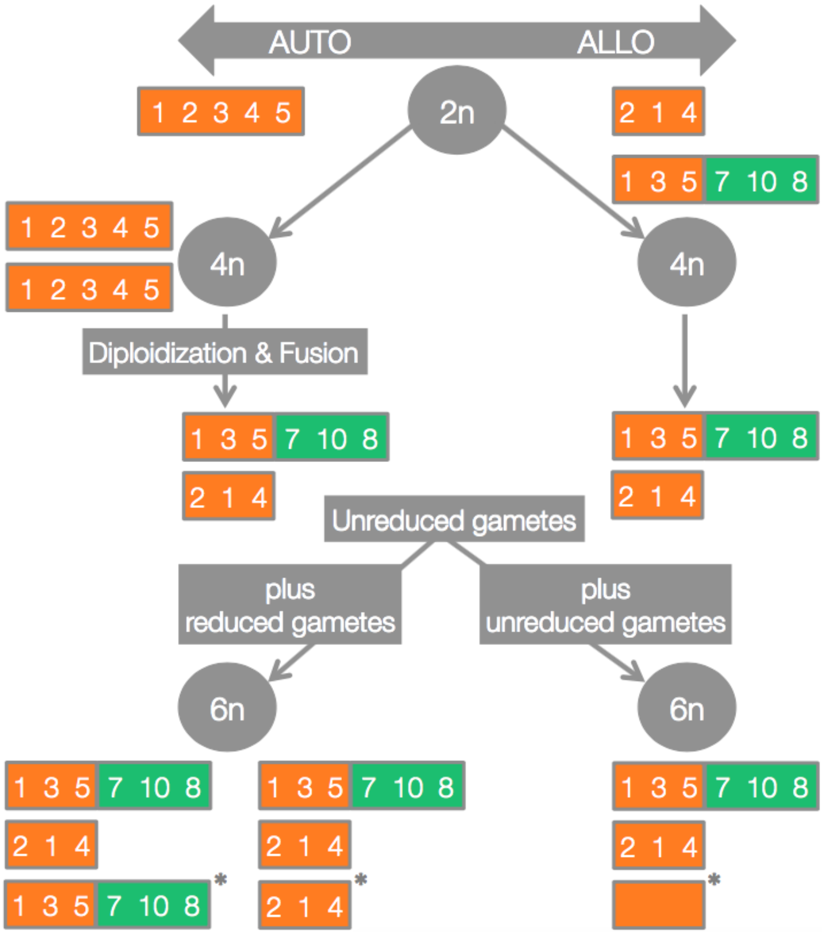


**Fig. S12: Expected segregation of an ancestral chromosomal fusion in grape under auto- versus allopolyploidy.** The overall scheme is similar to the one in Fig. S11 but additionally incorporates a chromosomal fusion. The fusion of "orange" and "green" chromosomes might have happened in the tetraploid eudicot genome (4n) following "autopolyploidy" **(left)** or in one of the diploid progenitor genomes (2n) before "allopolyploidy" **(right).** In the latter scenario, the diploid progenitor chromosomes do not only differ in their structure (i.e. fused versus nonfused), but also in their gene order on the corresponding homologous regions ("orange" rectangles). So, either mode of tetraploidy can give rise to structurally divergent chromosome pairs in the tetraploid genomes (4n). As in above, only hybridization with a nontetraploid-like genome can add the "outlier" third paralogous chromosome in grape **(bottom right)**.

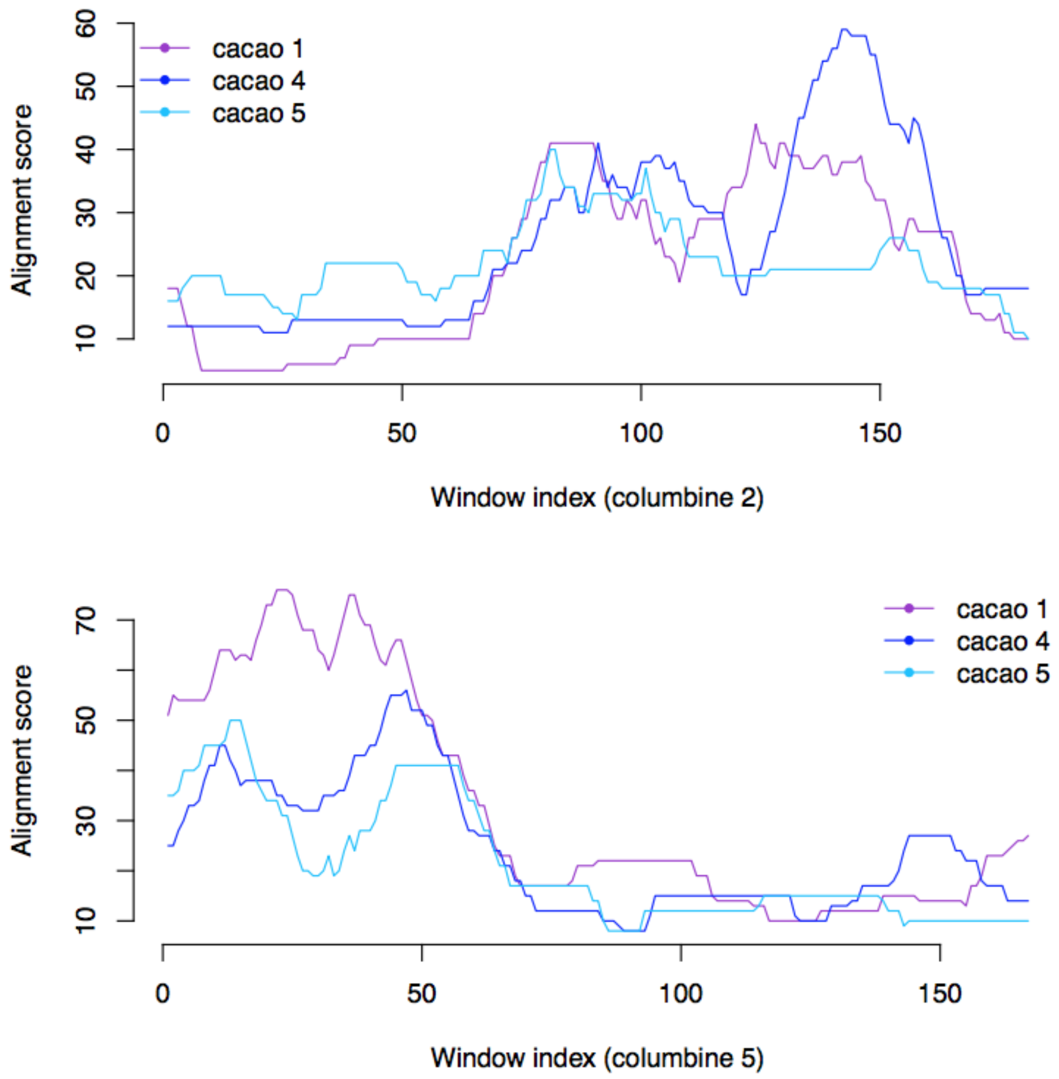


**Fig. S13: Examples of gene order similarity between the homologous regions of columbine and cacao.** For successive windows of genes within a given columbine chromosomal region, the best alignment score with respect to each of the three cacao chromosomes harboring homologous regions, is given. For example, columbine chromosomes 2 and 5 share a paralogous region homologous to cacao chromosomes 1, 4, and 5. The chromosome 2 region **(top panel)** appears to be most closely related to cacao chromosome 4, whereas its paralogous counterpart on chromosome 5 **(bottom panel)** appears to be most closely related to cacao chromosome 1. Note that the results presented here are shown for a window size of 30 genes but remain significant for all the window sizes tested (p=0-0.05).

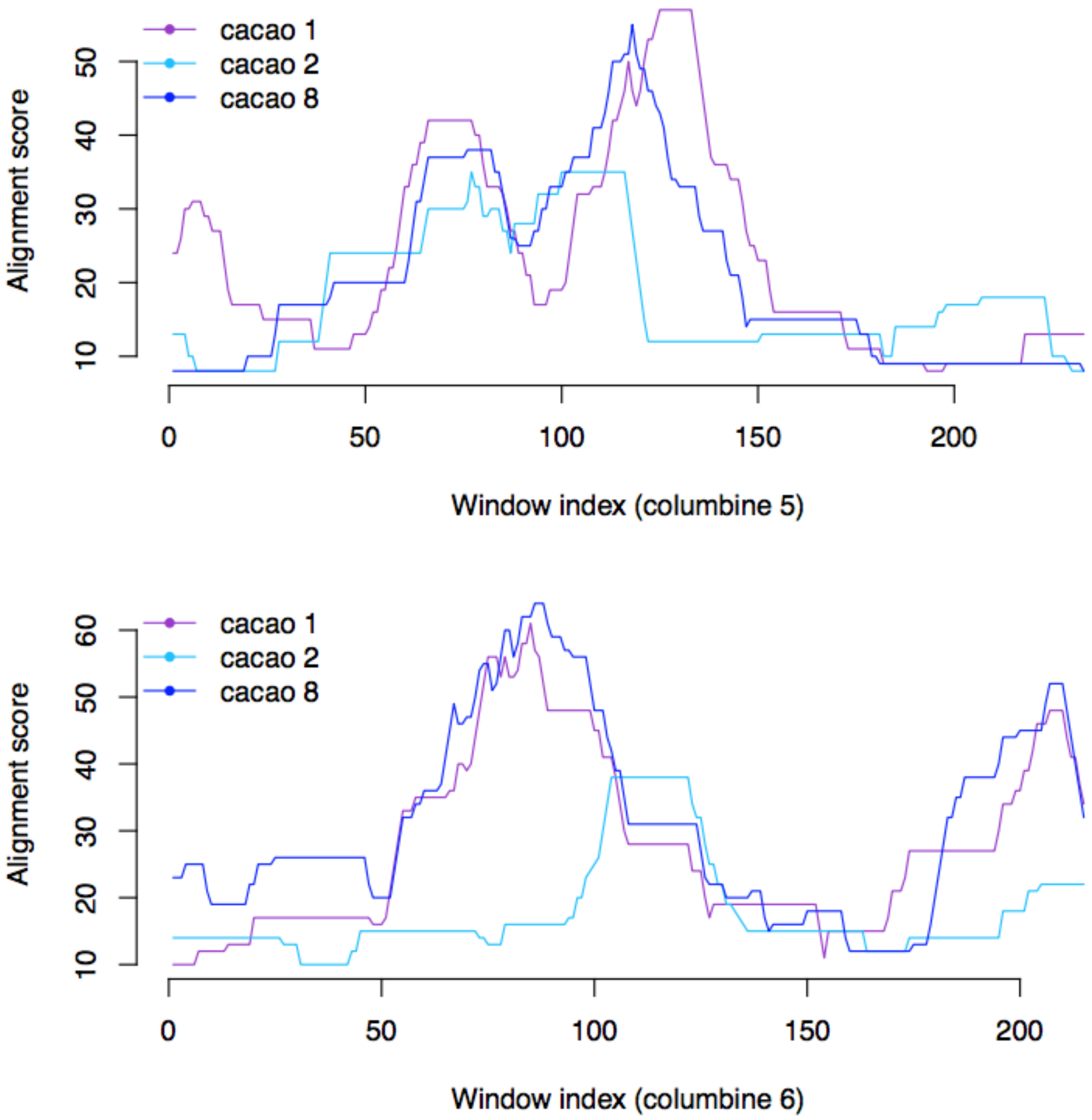


**Fig. S14: Examples of gene order similarity between the homologous regions of columbine and cacao.** For successive windows of genes within a given columbine chromosomal region, the best alignment score with respect to each of the three cacao chromosomes harboring homologous regions, is given. For example, columbine chromosomes 5 and 6 share a paralogous region homologous to cacao chromosomes 1, 2, and 8. The chromosome 5 region **(top panel)** appears to be most closely related to cacao chromosome 1, whereas its paralogous counterpart on chromosome 6 **(bottom panel)** appears to be most closely related to cacao chromosome 8. Note that the results presented here are shown for a window size of 35 genes but remain significant for all the window sizes tested (p=0-0.03).

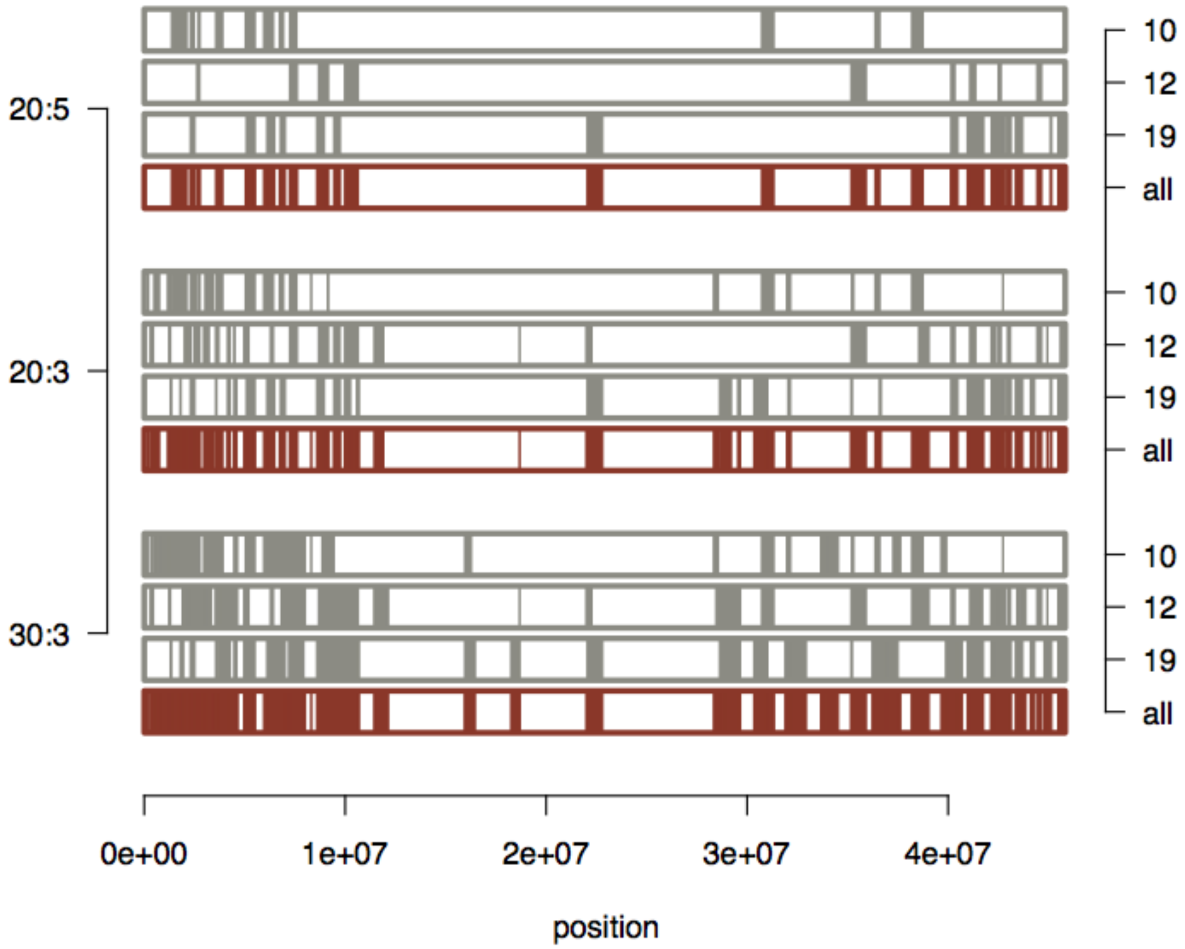


**Fig. S15: Synteny between columbine chromosome 4 and paralogous grape chromosomes 10, 12 and 19.** Results are shown for three different runs of DAGChainer. Parameter combination D:A (Y-axis on the left side) specifies the maximum genic distance between two matches (D) and the minimum number of aligned gene pairs (A) required to define a synteny block between columbine and grape. For a given parameter combination, columbine chromosome 4 is drawn four times in the order of its synteny to each of three the paralogous chromosomes and the overall synteny across all paralogous chromosomes (10, 12, 19, all; Y-axis on the right side). The overall synteny to all paralogs is highlighted in dark red.

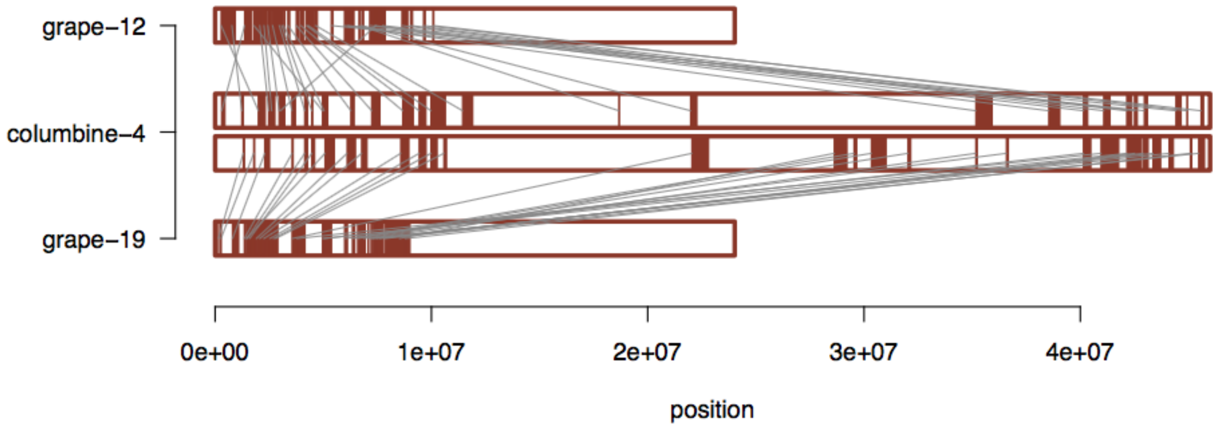


**Fig. S16: Synteny between between columbine chromosome 4 and grape chromosomes 12 and 19.** Note that this result is generated with the parameter combination D:A=20:3.

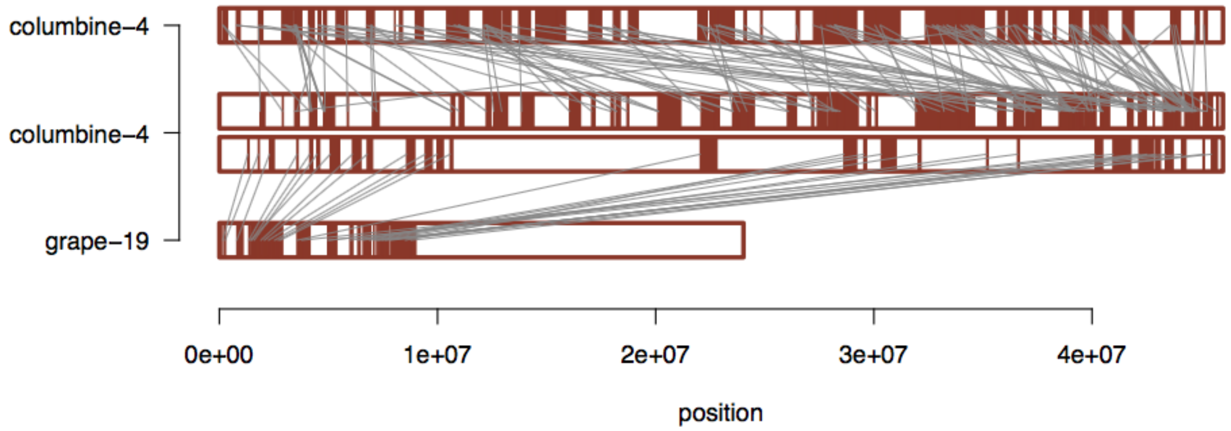


**Fig. S17: Zoom into syntenies of columbine chromosome 4 with itself and grape chromosome 19.** The results are generated with the parameter combination D:A=20:3 for both within and between genome synteny detection. Note that the most relaxed parameter combination (D:A=30:3) creates an even more complicated synteny relationship on chromosome 4 and thus is not shown here.

### Supplementary Data

1. columbine-to-columbine.synteny.blocks.and.genes.xlsx provides the list of synteny blocks in the columbine genome together with the paralogous gene pairs located in these blocks. Pink-filled cells correspond to the subset of blocks framed in Fig. S1.

2. columbine-to-grape.synteny.blocks.and.genes.xlsx provides the list of synteny blocks between columbine and grape chromosomes (sheet 1) and the homologous gene pairs located in these blocks (sheet 2). The former is used to show the synteny relationship between the two genomes (Figs. 3, S3 and S4) and also to color the columbine chromosomes as in Figs. 6 and S3.

3. grape-to-grape.synteny.blocks.and.genes.xlsx provides the list of synteny blocks in the grape genome (sheet 1) and the paralogous gene pairs located in these blocks (sheet 2). The former is used to color the grape chromosomes as in Fig. 6.

4. reconstruction.of.columbine.and.grape.chrs.xlsx provides the list of at least three genes consecutively matching between a pair of columbine and grape chromosomes and their paralogous counterparts on the remaining chromosomes of columbine and grape, respectively. In this table, zeros denote the absence of the latter and letters code for homologous genes as shown in Fig. S5.

5. homologous.genes.bet.columbine.and.grape.xlsx provides the list of homologous genes used for multiple protein sequence alignment.
